# Predicting Mutation-Induced Relative Protein-Ligand Binding Affinity Changes via Conformational Sampling and Diversity Integration with Subsampled Alphafold2 in Few-Shot Learning

**DOI:** 10.1101/2025.02.09.637298

**Authors:** Liangxu Xie, Xiaohua Lu, Dawei Zhang, Lei Xu, Mingjun Yang, Young-Seob Jeong, Shan Chang, Xiaojun Xu

**Author notes:** Correspondence: S.C.; X.X.

## Abstract

Predicting the impact of mutations on protein-ligand binding affinity is crucial in drug discovery, particularly in addressing drug resistance and repurposing existing drugs. Current structure-based methods are constrained by their dependence on known protein-ligand complex structures. The absence of structural data for mutated proteins, along with the potential of mutations to modify protein conformational states, brings challenges for reliable predictions. The downstream application of conformational sampling of Alphafold, such as the prediction of binding free energy changes induced by mutation, has not been fully explored. To tackle the challenge, we propose a few-shot learning approach that integrates AlphaFold2 subsampling, ensemble docking, and Siamese learning to predict the relative binding affinity changes induced by mutations. We validate our approach for predicting binding affinity changes in 31 variants of the mutated Abelson tyrosine kinase (ABL). The conformation ensembles for each ABL variant are generated using AlphaFold2 subsampling. The most likely conformational states are selected by Gaussian fitting to construct the pairwise structural inputs and compute relative binding energies. Then the relative binding affinities are derived by averaging the values predicted by the Siamese learning model across the constructed conformational pairs. By benchmarking on the Tyrosine kinase inhibitors (TKI) dataset and the direct binding affinity using the refined set of PDBbind database, our proposed approach achieves an increased Spearman correlation coefficient for five out of six TKI molecules across 31 mutants when estimating relative binding affinities. Although the proposed method is validated on the ABL, it can be transferred and applied to other drug-target interaction predictions when the conformational flexibility of proteins needs to be considered. We anticipate the approach will be valuable for predicting binding energy changes induced by mutations in proteins or ligands, facilitating drug discovery efforts.

## Introduction

Proteins are responsible for life-essential specific functions such as cellular structural support, immune protection, enzyme catalysis, cell signaling to transcription, and translation functions ^1-5^. These functions are closely related to structural changes in the proteins. The experimentally determined three-dimensional structures remain limited due to the complexity and expense of techniques like X-ray crystallography, nuclear magnetic resonance, and electron cryo-microscopy (cryo-EM). Consequently, prediction of protein structures through computational methods has become crucial ^6-8^. Deep learning techniques, notably AlphaFold2 (AF2), have revolutionized the field of protein structure prediction. AF2 provides prediction accuracy close to experimentally determined methods ^9^. Following the success of AF2, many predictors have been generated such as RoseTTAFold ^10^, ColabFold, OpenFold ^11^; ESMfold ^12^, RGN2 ^13^, and OmegaFold ^14^. The success of AF2 brings new possibilities for many downstream explorations, such as analyzing missense variants in genome sequences ^15^, predicting protein-ligand molecule interaction in drug discovery ^16,17^, protein engineering in synthetic biology ^18,19^, and metallic pathway analysis for biological systems ^20^.

While it effectively predicts static protein structures, the default AF2 has limited capability in capturing the dynamic behavior and conformational changes of proteins. Proteins are not static and undergo conformational changes upon binding of a specific ligand, whereby residues in the pocket adjust their conformation to optimize their interaction with the ligand ^21,22^. Significant efforts have been dedicated to refining AF2 to capture different conformations. Recent advancements in protein design have facilitated the detailed characterization of protein dynamics, enabling the identification of both ground and excited states ^23^. The AF2 model constructs protein 3D structures by using Multiple Sequence Alignment (MSA) as inputs ^24^. Since MSA encodes coevolutionary information, manipulating MSA offers a viable way to alter the conformations by controlling coevolutionary information. Several successful methods have been reported for predicting conformational ensembles of proteins, such as AF-Cluster via sequence clustering ^25^, subsampling MSA ^26^, energetic frustration analysis of MSA ^27^, SPEECH_AF via in silico mutagenesis of MSA ^28^, randomized alanine sequence scanning ^29^, etc. The recent successful cases have applied AF2 to directly predict the relative conformation populations by subsampling the comparison of multiple sequences ^26,30^. Additionally, analysis of MSA has also been used to identify patterns of relevant mutations within protein families ^31,32^, predict interacting residues based on sequences ^33,34^, and construct structure prediction models guided by energy-based processes ^35,36^.

After predicting the structures of proteins, binding affinity can be predicted for assessing the strength and stability of small molecules binding to protein targets. With advances in databases and computational power, predictions can be made using physics-based methods and deep learning methods ^37-40^. However, the binding sites of proteins in different conformational states can exhibit significant variations, which may affect the binding mode. Consequently, the challenge has changed from merely determining whether a protein structure is available to appropriately selecting an optimal structure for accurate modeling. To avoid bias toward a single protein conformation and to implicitly account for protein flexibility, an ensemble of representative structures is used to dock candidate ligands ^41^. Incorporating protein flexibility is essential for accurate binding affinity prediction, as exemplified by studies on Abelson tyrosine kinase (ABL) ^42^. Although Tyrosine kinase inhibitors (TKIs) like imatinib have revolutionized the treatment of chronic myeloid leukemia (CML) ^43^, drug resistance emerges from mutations that alter the conformation, thereby affecting the drug binding mode and binding affinity ^44^. The limited number of crystal structures of kinase mutants poses a limitation to the accuracy of both physics-based methods and machine/deep learning methods ^45^.

Evaluating the changes in binding affinity caused by mutations is crucial for understanding essential biological processes. Despite considerable endeavors, accurately predicting the impact of mutations on binding affinity remains challenging ^46-48^. Mutation-induced structural changes bring two major challenges for the traditional computational methods: determining the precise structural changes of proteins, and selecting the most representative structure from conformation ensembles. A real-world scenario in drug development involves understanding the resistance profiles of certain drugs against various mutated structures ^49^. For example, mutations in ABL alter the relative population of the active state and inactivate state, which can be selectively targeted by different types of inhibitors ^50^. Several states of ABL are structurally similar to known crystal structures capable of binding a variety of inhibitors ^51^. Consequently, the conformation ensemble of kinase proteins should be incorporated into the prediction of binding free energy changes induced by mutations. The accuracy of AF2 extends to the resolution of individual amino acids, enabling precise predictions of protein structures at an atomic level ^15,52^. AF2 has learned an approximate biophysical energy function that can accurately assess the quality of mutated protein structures ^53^. Furthermore, subsampled AF2 is capable of predicting the conformational landscape and the relative populations of conformations at single mutation resolution ^26^. In this study, we propose an integrated approach to address the aforementioned limitations by streamlining AF2 subsampling, ensemble docking, and Siamese learning. To incorporate structural diversity, we employ data augmentation by pairing the reference structures with those generated structures from subsampled AF2. We benchmark our proposal by investigating drug resistance in ABL kinase. To efficiently process the structural features, we propose an STGNet (Structure and graph-aware Siamese network) model in the Siamese architecture because Siamese network has been used to predict relative binding affinity for the static structures ^54^. In few-shot learning, our approach demonstrates higher ranking accuracy in predicting binding free energy changes induced by mutations.

## Results

### Construction of augmented pairwise dataset and deep learning architecture

Predicting relative properties has significant applications in drug discovery, for example, changes in absorption, distribution, metabolism, excretion, and toxicity (ADMET) during drug lead optimization ^55,56^, as well as in assessing relative binding affinities between congeneric ligands ^57^. Various models have been proposed for this purpose as illustrated in **Fig. 1**. The relative properties can be computed using a standard neural network, the delta model ^56^, or the Siamese model ^54^. The Siamese architecture is designed to compare two inputs by generating embeddings, thus capturing subtle distinctions between similar inputs ^54^. The pairwise Siamese network further expands on this concept by integrating pairs of representations and calculating average values across various inputs ^55^.

**Fig. 1:**
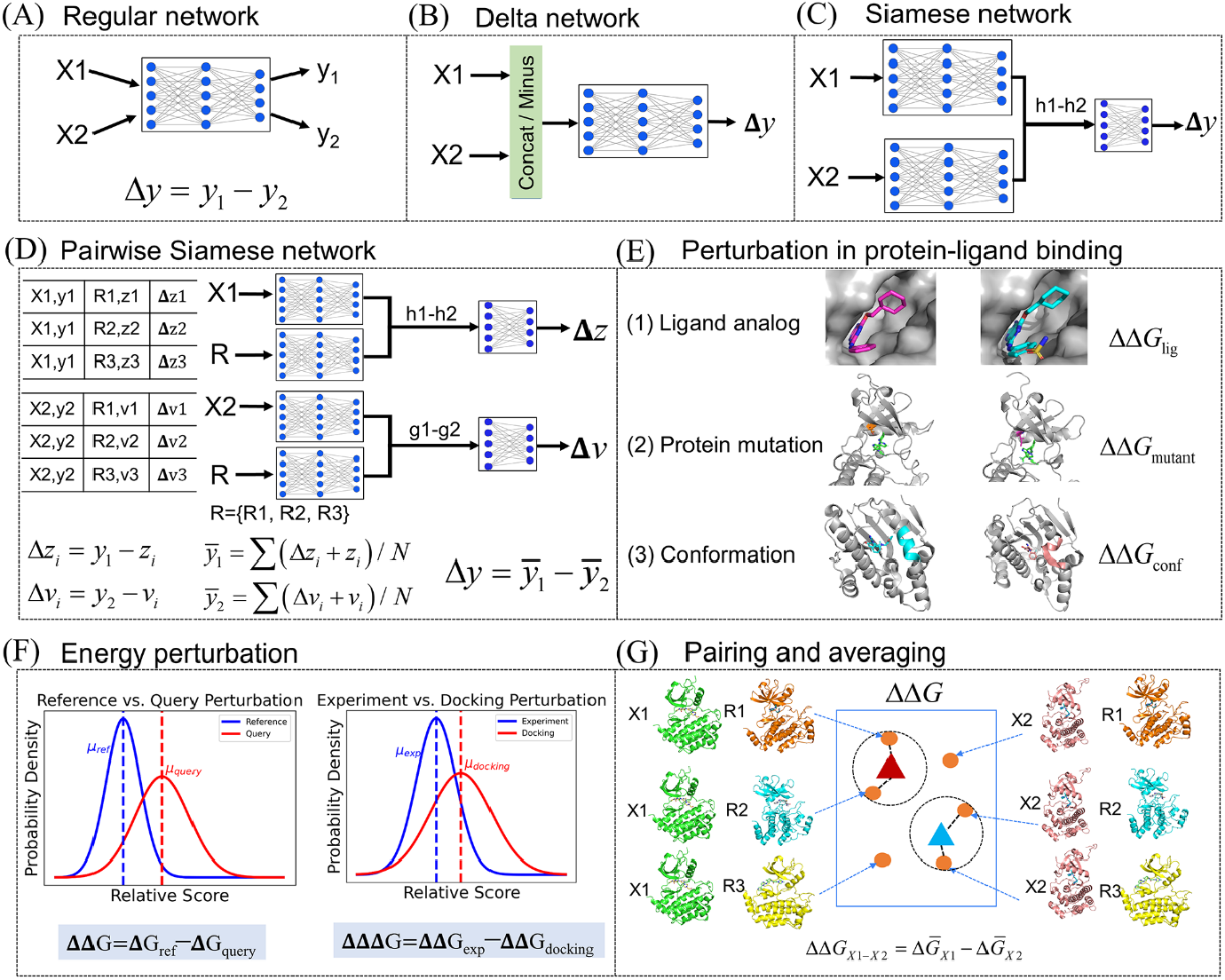
Models for relative property prediction. (A) Regular neural network. (B) Delta neural network. (C) Siamese network. (D) Pairwise Siamese network. (E) Perturbation in protein-ligand binding includes variations of ligand analogs, protein mutation, and conformations. (F) Two types of energy perturbation. (G) Pairing and averaging with the most populated states when estimating the relative binding free energy. X1 and X2 represent the input, and y_1_ and y_2_ are the output. R1, R2, and R3 represent the reference states.

In addition to deep learning methods, free energy perturbation (FEP) is a widely employed physics-based method for predicting relative binding affinity ^58^, which quantifies energy changes between a reference and a target state. In our study, we broaden the scope of perturbation analysis by transitioning from absolute value comparisons to evaluating relative values, as illustrated in **Fig. 1F**. Specifically, we focus on analyzing the relative changes in binding energy induced by mutations. To achieve this, we leverage deep learning techniques to establish a mapping between relative scores and the corresponding relative binding energies associated with mutations. To enhance the reliability of our predictions, we also implement a pairing and averaging process as shown in **Fig. 1G**. This method will be useful for computing relative binding affinities in protein-ligand complexes that are caused by ligand structure transformation, protein mutations, and conformational variations during interactions.

The augmented dataset is constructed by using AF2 subsampling and ensemble docking (refer to the Method section for details in supporting information). In the workflow illustrated in **Fig. 2**, we selected 31 mutants of ABL kinase and the 6 TKI molecules to construct our dataset. The objective is to construct reference-generated conformation pairs for the specified mutants. The reference values are derived from the TKI dataset ^59^. The generated values are predicted from subsampling AF2 for 31 mutants and wild-type of ABL kinase. The relative docking score ΔΔ*G*_dock_ is computed by comparing the docking scores of mutants to those of the wild type.

**Fig. 2:**
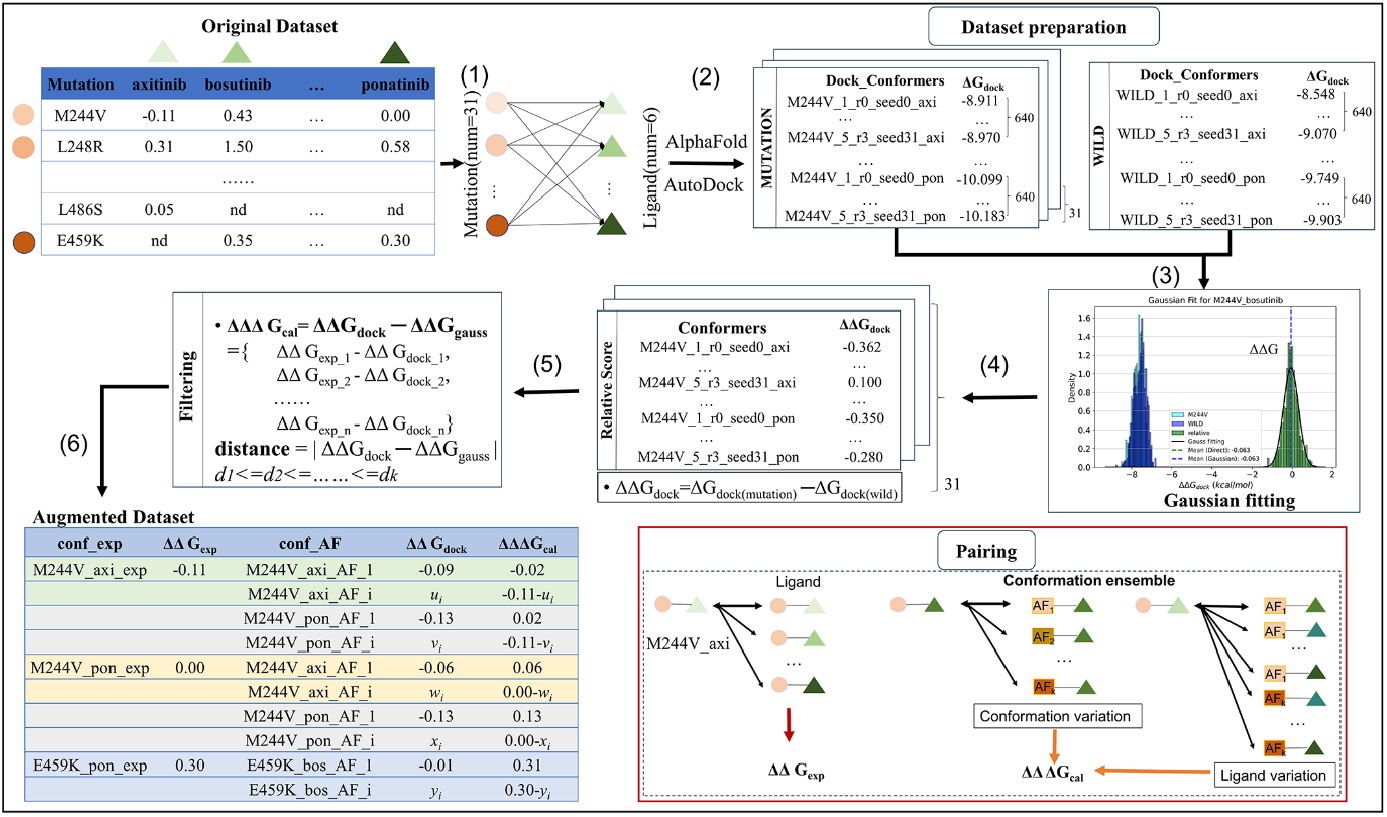
Pipeline of construction of the augmented pairwise dataset. The workflow includes the creation of initial mutant-ligand data pairs, conformational sampling, ensemble molecular docking, relative binding energy calculation, data filtering, and construction of the final dataset. The red bounding box highlights the pairing between the reference state and the query states, encompassing the conformation variation and ligand variation.

From the distributions of relative docking scores, we identified the most probable conformations. The selected relative docking scores are then paired with the experimental values. The relative binding affinity for each experimental value will be augmented to a specified number of data points. Then, we can obtain the relative binding free energy difference (referred to as ΔΔΔ*G*_pred_) by subtracting the reference value from the relative docking score.

By specifying the mutation type, this approach emphasizes differences in conformation and variations in bound ligands. The key advantage of using the relative binding affinity difference is that it allows Siamese learning to focus on the difference between the reference state and the generated conformational states. Experimental-level binding energy accuracy can be achieved through the equation ΔΔ*G*_exp_ = ΔΔ*G*_dock_ + ΔΔΔ*G*_pred_ . Other computational details can be found in the Methods section.

Data augmentation has been demonstrated to enhance the prediction of binding affinities, offering improved accuracy and robustness ^39^. The impact of multi-conformation data on the predictive ability of the model is evaluated by systematically varying the number of conformations, specifically using *k* values of 1, 5, 50, and 100, respectively. This methodology expands the dataset through *k*-fold augmentation by pairing the experimental data with the generated data, incorporating structural diversity for each protein-ligand pair via conformation variation and ligand variation. After obtaining the augmented dataset, the dataset is partitioned into training and test sets according to the names of different TKIs for cross-validation. This splitting method reflects real-world scenarios where we have established knowledge of protein mutations and their associated changes in binding free energy. By leveraging this prior knowledge, we can predict the relative binding free energy of a new drug molecule across all mutations. Further details are described in the Methods section and Fig. S1 in supporting information. All molecules exhibit low similarity as shown in Fig. S2, which ensures that pairing with the experimental values does not result in information leakage. Similar pairwise approaches have been employed to predict relative binding free energies ^54,57,60^. Our approach differs from existing work in two key respects. First, previous research, such as pairwise binding comparison network (PBCNet), has focused on predicting changes in binding energy due to variations in ligands. In contrast, our method extends PBCNet by specifically addressing the prediction of binding energy changes caused by protein mutations. Secondly, we introduce the concept of using the difference in relative binding free energy as the prediction target, rather than the relative binding free energies themselves. This modification enables the Siamese learning model to more accurately capture the differences among various conformation states.

To harness the structural information, the input for our Siamese learning includes contact maps of each conformation, the molecular graph of ligand, and the molecular graph of protein-ligand interaction as shown in **Fig. 3**. The molecular graph has been validated in our previous work ^61^. The protein contact map with BLOSUM62 matrix scores ^62^ can reflect the spatial contacts between CA atoms of each residue in the three-dimensional (3D) structure of a protein. The ligand graph contains the chemical properties within the molecule. The protein-ligand interaction graph considers covalent interaction between intramolecular and non-covalent atomic-level interaction information of intermolecular. To learn the structural features, we constructed the STGNet (Structure and graph-aware Siamese network) model to learn the structural information using the Message Passing Neural Networks (MPNN) network ^63^, and to learn interaction information using the Geometric Interaction Graph Neural Network (GINN) network ^64^. The protein contact map and ligand graph are learned using MPNN networks in the Siamese architecture, which helps to capture the 3D folding structures of the protein as well as the internal chemical properties of the ligand. GIGN can learn the binding patterns between proteins and ligands, by combining the properties of the nodes and the 3D coordinate information. By using MPNN and GIGN together, the model can focus on intramolecular and intermolecular features, respectively. The experimental values and the paired AF2-generated data will be fed into the shared Siamese Network. Therefore, the network can better learn the distinctions between the reference state and the alternative conformational states, thereby enhancing the prediction of relative binding affinity differences.

**Fig. 3:**
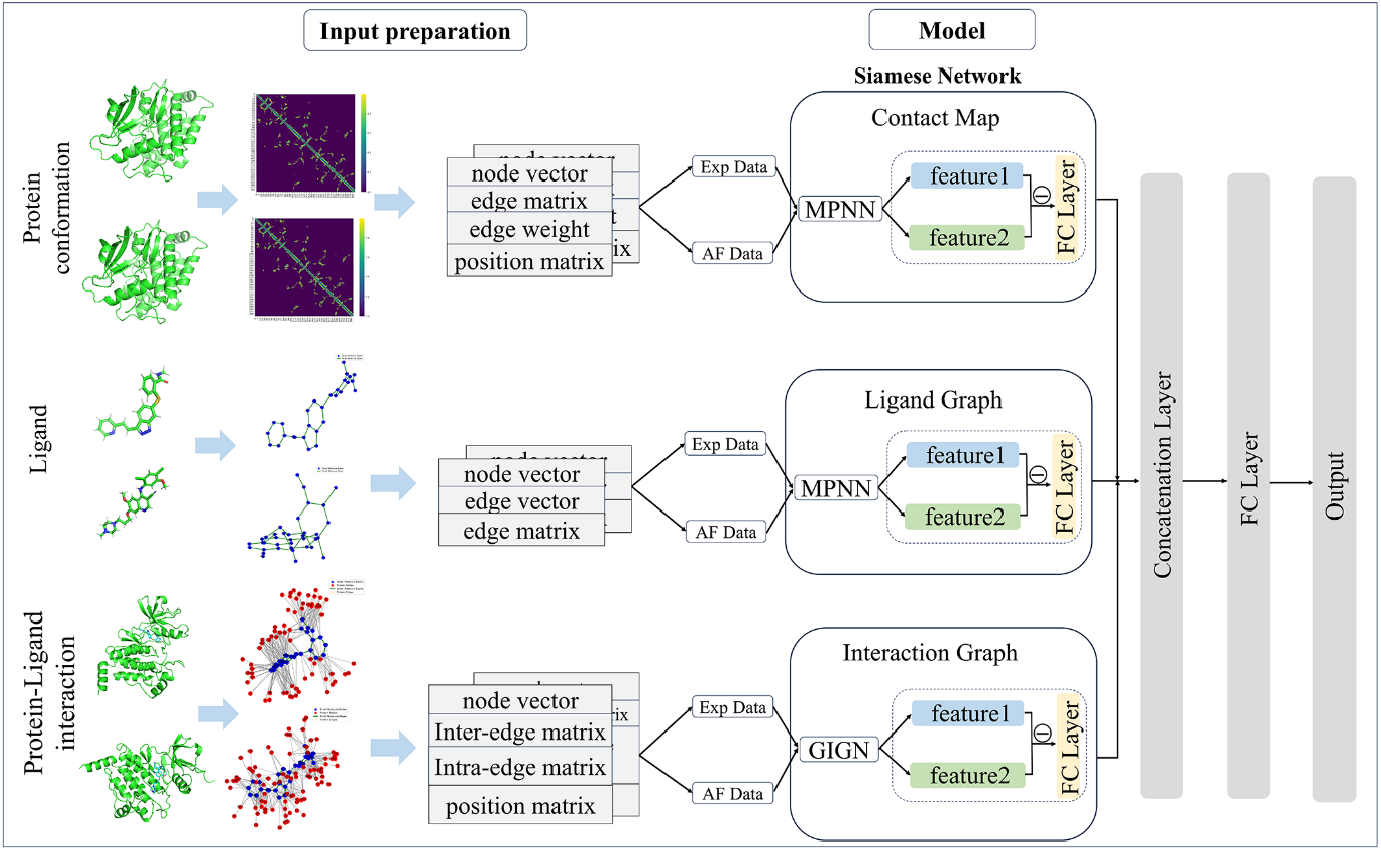
Architecture of the STGNet model. The reference state and the AF2-generated state are processed by the Siamese network to capture the feature difference. Protein contact maps and ligand graphs are processed by MPNN, and protein-ligand interaction graphs are processed by GIGN. Ligands show as blue nodes and edges, and proteins show as red nodes and edges in molecular graphs.

### Conformation sampling for ABL kinase using subsampled AF2

Conformational changes in ABL kinase have important implications for its function and drug resistance ^65^. Mutations induce obvious structural changes in the P-loop and A-loop region, thereby modulating the binding affinity of the drug to ABL kinase by interfering with the stability of specific conformations (e.g., DFG-in and DFG-out) ^45,66^. Structural analyses comparing type I (DFG-in) and type II (DFG-out) inhibitors reveal that these two classes of drugs exhibit different sensitivities to conformational changes in ABL kinase. **Fig. 4** demonstrates three conformational states of ABL kinase, including the active state, inactive state I_1_, and inactive state I_2_. In the active state, P-Loop and A-Loop are in the open state and αC-Helix is in an inward position. Inactive state I_2_ shows obvious structural changes with active state, with the αC-Helix shifting from the inward position to the outward position, the P-Loop becoming closed, and the A-Loop partially folding. The inactive state I_1_ displays relatively minor conformational changes compared to the active state. Mutations may indirectly reduce the availability of drug-binding sites by stabilizing a certain conformation state and altering the percentage of the active state of the kinase ^67^.

**Fig. 4:**
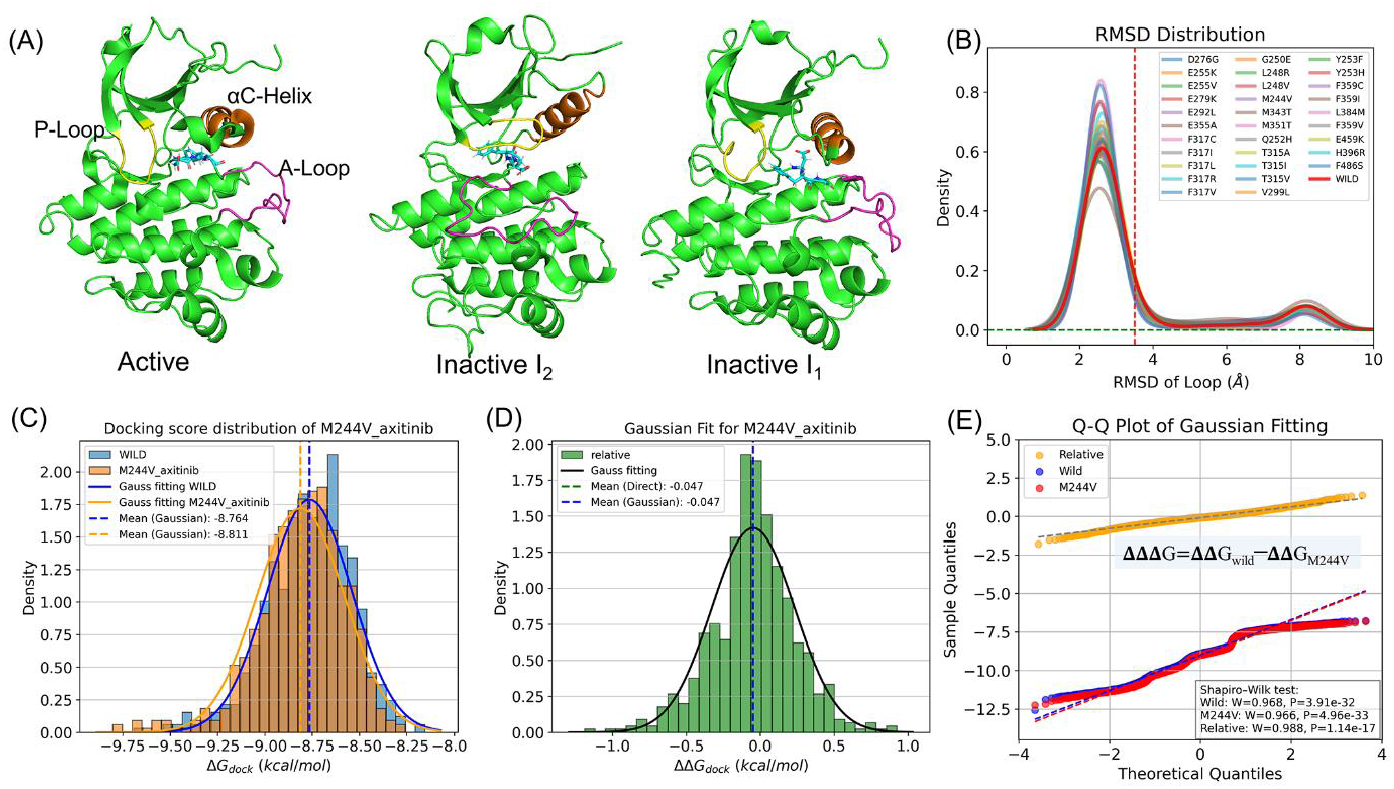
Mutation effect on the conformation of ABL kinase and the drug binding affinity. (A) ABL kinase conformation in active state (PDB ID: 6XR6) and two inactive conformational states I_1_ and I_2_ (PDB ID:6XR7, 6XRG). (B) RMSD distribution of the loop regions (A-loop, P-loop, and C-helix) for wild type and 31 mutants. (C) Distribution of AutoDock Vina docking score of the wild and mutant (M244V) of ABL kinase. (D) Density distribution plot based on the relative binding energy obtained from the mutant (M244V) and the wild type. (E) The Quantile-Quantile Plot for the docking score and the relative docking score for the mutant (M244V) and the wild type. More information can be found in Fig. S3-S8.

The previous study also presents a conformational dynamic equilibrium between the active and two inactive states (I_1_ and I_2_) of 8 mutants of ABL kinase achieved by subsampling MSA^26^. We adopted the same procedure of subsampling MSA and obtained the conformation ensemble for all 31 mutants. The deviation in conformation distributions of the mutants relative to the crystal structure is quantified by computing root mean square deviation (RMSD). **Fig. 4B** depicts the RMSD distribution densities of the loop regions of 31 mutants and the wild-type (WILD). The primary peaks of the RMSD densities for the wild-type are concentrated at RMSD values less than 3.5 Å, which is consistent with previous study ^26^. For wild type, the percentage of active state is 83%, which is close to the experimental value of 88%. The different distributions suggest that the mutation alters the conformational balance of the ABL kinase. The relative population of other mutants is summarized in Table S1.

The distribution of docking score for mutant-ligand (M244V_axitinib) and the wild-type (WILD_axitinib) are shown in **Fig. 4C-D**. The Gaussian fitting indicates the binding free energy distribution follows the principle of maximum likelihood. **Fig. 4E** confirms the Gaussian distribution of relative binding free energy distributions ΔΔ*G*_dock_ of the M244V mutant and the wild-type of ABL kinase, as validated by the Sharipo-Wilk test. Similar Gaussian distributions are observed for other muations as shown in Figs. S3-S8. The docking distribution can be interpreted as a perturbation of the experimental distribution. Consequently, our objective is to determine the relative energy changes induced by mutation. Based on the principle of maximum likelihood, we select the *k* conformations and their relative binding affinities, ensuring that the deviation of these affinities from the Gaussian mean value is minimized. The proposed filtering approach provides a robust statistical framework for identifying the most populated states, rather than simply prioritizing the highest-scoring conformation.

### Improved predictive performance over physics-based methods (docking and FEP)

Prediction of relative binding free energy delivers import value in drug discovery applications. However, accurate prediction is limited due to technical and practical challenges in physics-based modeling and statistical mechanics methods ^68^. The molecular docking and free energy perturbation (FEP+) were compared to our model. The predicted relative binding affinity and the experimental value are shown in **Fig. 5**. Molecular docking can provide a quick ranking, but it fails to reproduce the relative binding free energy between them. FEP+ can reproduce the relative binding affinity at the chemical accuracy, but it requires high computational resources. For our method, we provide the relative binding affinity for all experiment-AF2 pairs, then we compute the statistical average for the required protein-ligand pair, which enhances the stability of the prediction.

**Fig. 5:**
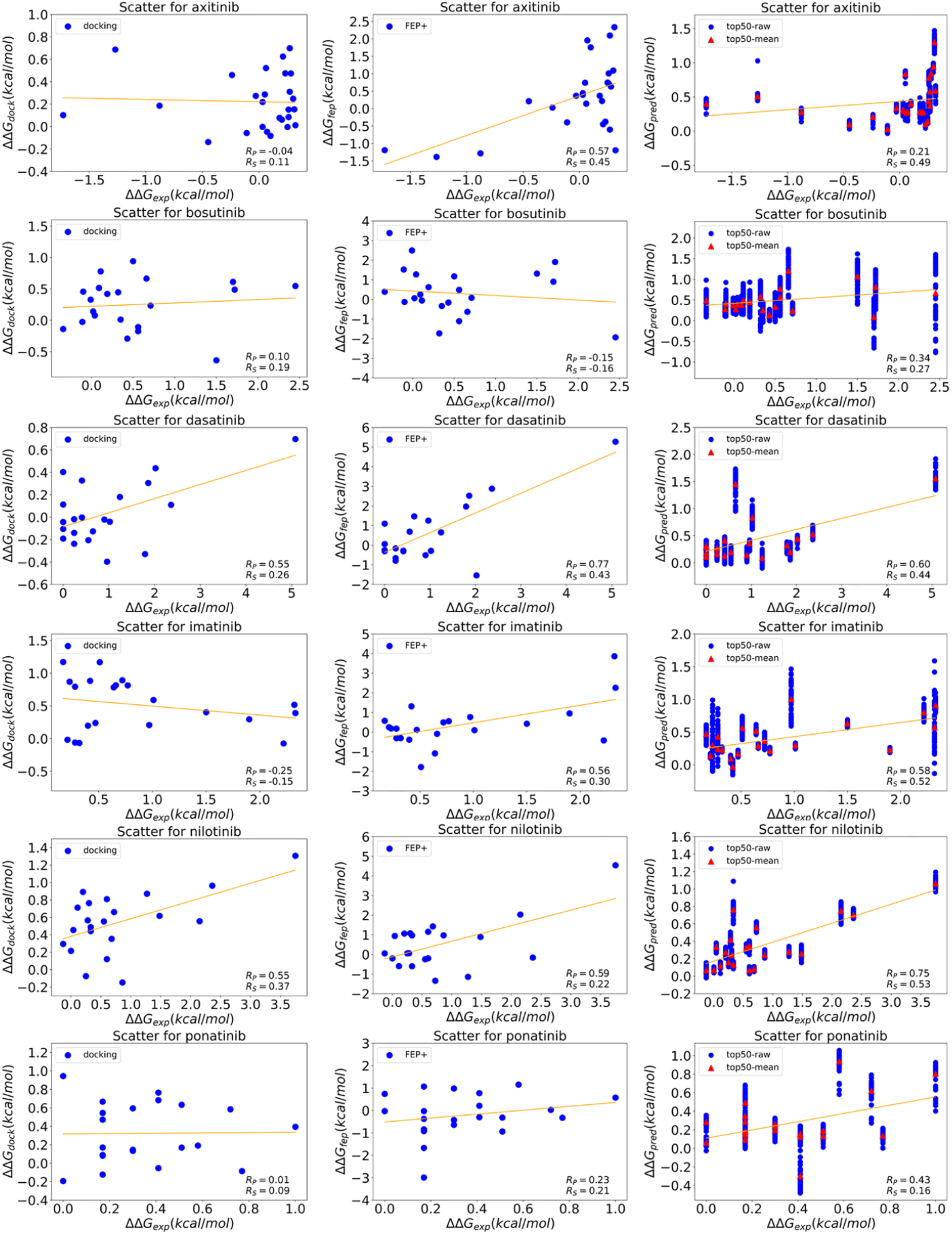
Comparison of experimental and predicted relative binding energies for six TKI molecules. The predictions were generated using molecular docking, FEP+, and STGNet (denoted as top50-raw for the raw prediction and top50-mean for the average over the 50 data points derived from pairs sharing the same reference state). Each blue sphere represents one prediction and the red triangle represents the average value.

The performance of the models was evaluated by calculating RMSE, MAE, Pearson, and Spearman coefficients, which are summarized in **Fig. 6** and Table S2-S5. It can be seen that molecular docking has limited performance for protein-ligand relative binding free energy calculation, as indicated by the low Pearson and Spearman coefficients. The FEP+ results reported by Hauser et al ^59^, are markedly superior to those obtained from molecular docking, as evidenced by higher Pearson and Spearman coefficients for the axitinib, dasatinib, imatinib, and ponatinib significantly higher than those of molecular docking. The STGNet, employing 50-fold data augmentation (denoted as top50), demonstrates enhanced performance in terms of RMSE, MAE, and Pearson coefficients, obtaining the best values for four types of TKIs: bosutinib, imatinib, nilotinib, and ponatinib. For the computational accuracy, our model cannot competed with The FEP+ results. For the ranking ability, our method outperforms both molecular docking and FEP+ methods, as evidenced by the highest Spearman correlation coefficients obtained for five out of six TKIs: axitinib, bosutinib, dasatinib, imatinib, and nilotinib, highlighting its enhanced ranking performance. By introducing a multi-conformational free energy distribution filtering strategy and applying Siamese learning, our method significantly improves the accuracy in predicting the relative binding free energy changes induced by mutation.

**Fig. 6:**
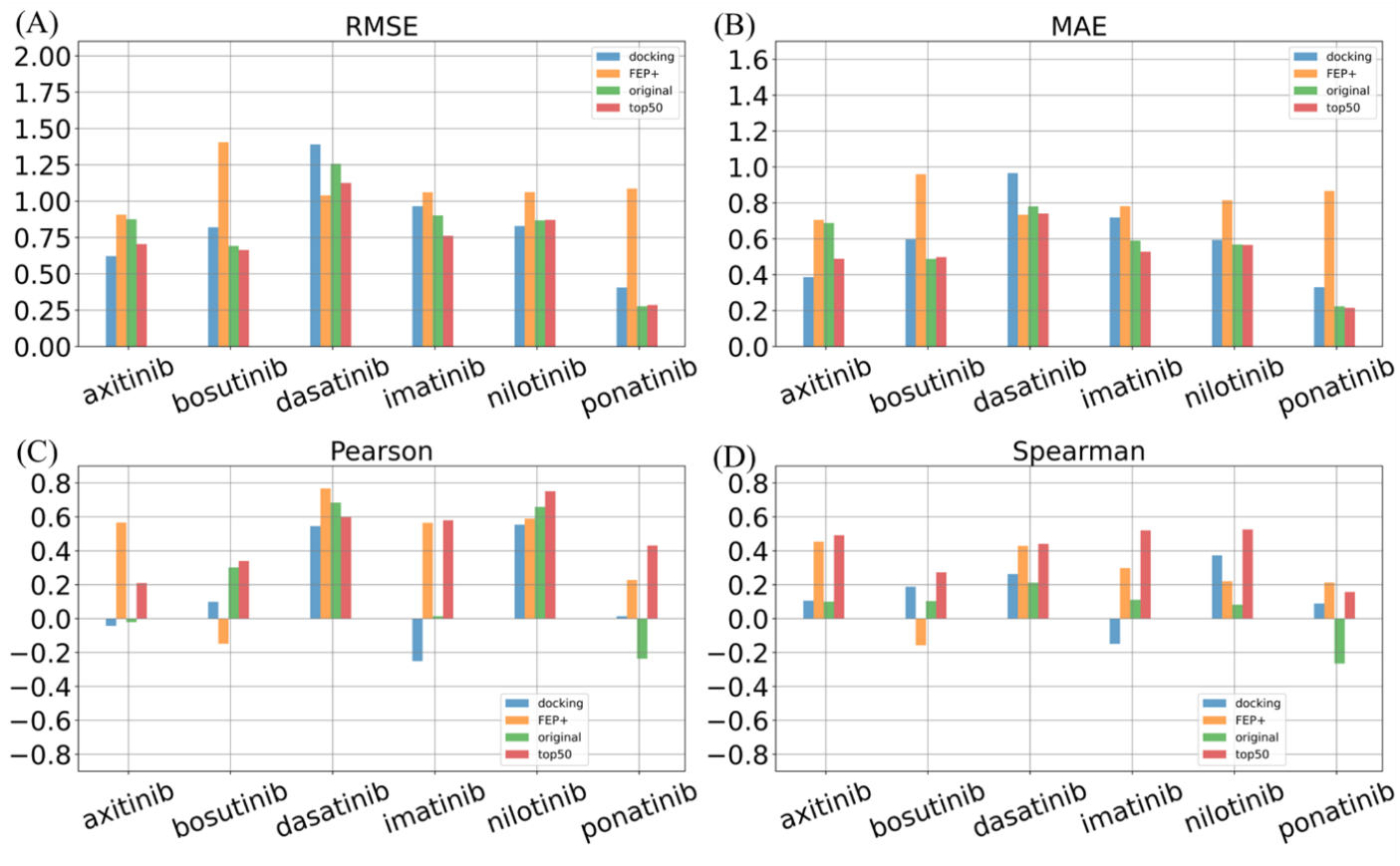
Evaluation metrics of four different methods. Performance is evaluated by RMSE, MAE, Pearson coefficients, and Spearman coefficients for docking, FEP+, STGNet using the original TKI dataset (denoted as original, which represents that no conformational flexibity is included), and STGNet using the top 50 samples (denoted as top50).

### Relative value prediction outperforms absolute value prediction

We compare the data augmentation method with direct absolute prediction of binding energy using the refined set of PDBbind after excluding the TKI-related data. The model for the regular neural network, illustrated in Fig. S9, was trained using a comparable network architecture except for omitting the Siamese network and the pairwise input. The model predicts the absolute value of the binding free energy. As shown in **Fig. 7A**, the regular model achieves satisfactory performance in predicting absolute binding affinities, with strong correlations (Pearson and Spearman coefficients are 0.90, and 0.90 for the hold-out set, and for the CASF-2016 set). However, the model exhibits limitations in predicting both absolute and relative binding energies for ABL mutants, as illustrated in **Fig. 7B and 7C**. Specifically, it fails to accurately capture the effects of single mutations in proteins. The Spearman coefficients in **Fig. 7D** support this observation, indicating the regular model’s inability to reliably predict relative binding energy changes induced by single mutation in proteins.

**Fig. 7:**
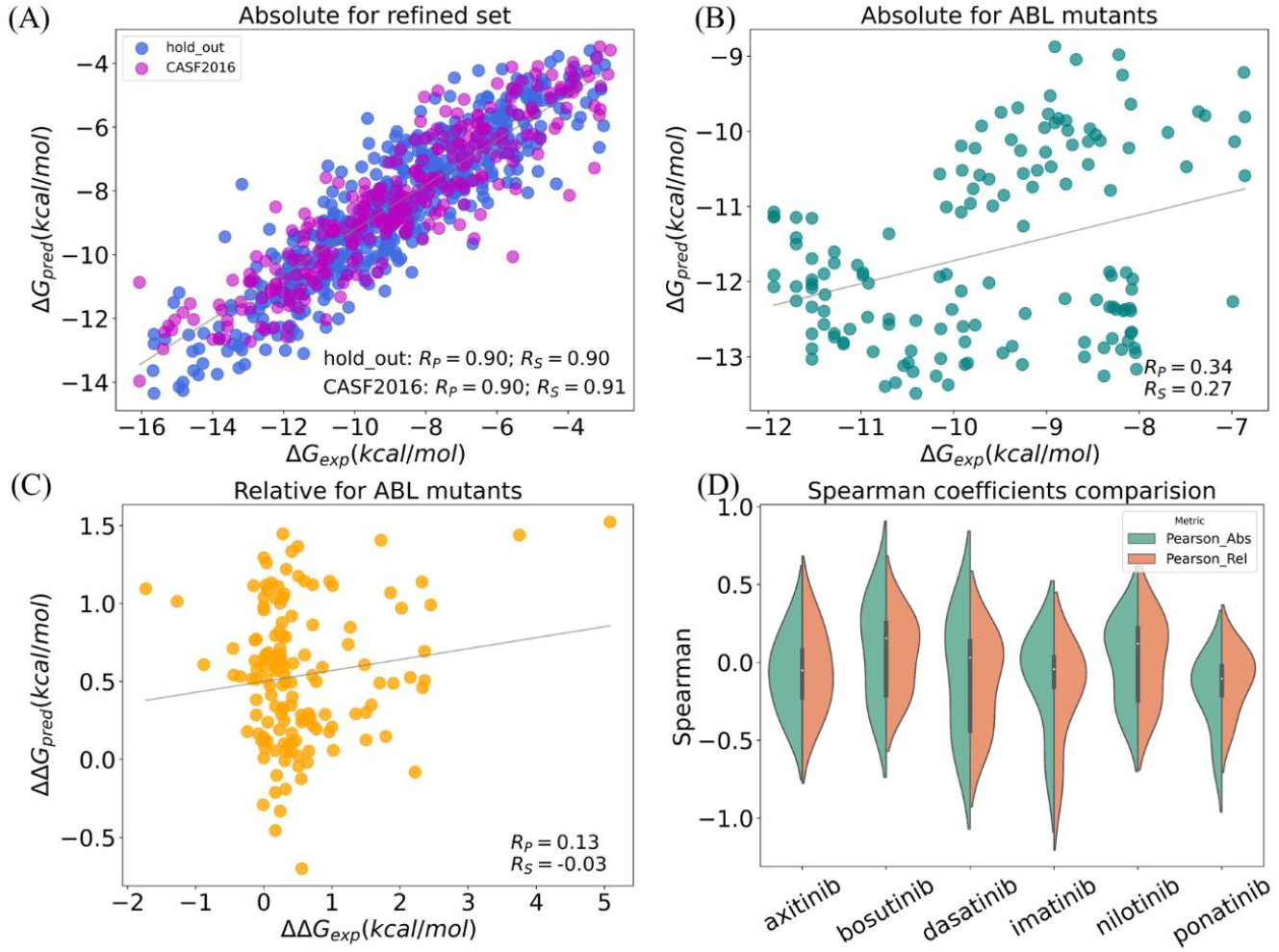
Performance of regular network in the prediction of absolute binding energy. (A) Correlation between the absolute values of the prediction and experimental values in the CASF-2016 set. (B) Correlation between the absolute values of prediction and experimental values in the TKI set. (C) Correlation between the relative values of prediction and experimental values in the TKI set. (D) Spearman correlation coefficients for both absolute values and the relative values of predictions compared to experimental values in five-fold validation.

We compare the result of the regular model with our STGNet model. As can be seen in **Fig. 8**, the RMSE and MAE of STGNet using the different numbers of samples (5, 50, and 100 are denoted as top5, top50, and top100) were lower than those of the refined dataset for five out of six molecules, including axitinib, bosutinib, dasatinib, nilotinib, and ponatinib. STGNet presents higher Spearman coefficients than that of the refined set. Combining experimental and AF2 data, the STGNet method is better able to capture the dynamic changes of proteins in different conformations and the effects of mutations on these changes. By combining the results from **Fig. 7** and **Fig. 8**, we demonstrate that our approach significantly enhances the ability of the STGNet model to recognize and interpret structural and energetic perturbations induced by single amino acid substitutions. Overall, our method enhances mutation effect prediction by integrating multiple conformations, applying a rational filtering strategy, and Siamese learning with the STGNet model.

**Fig. 8:**
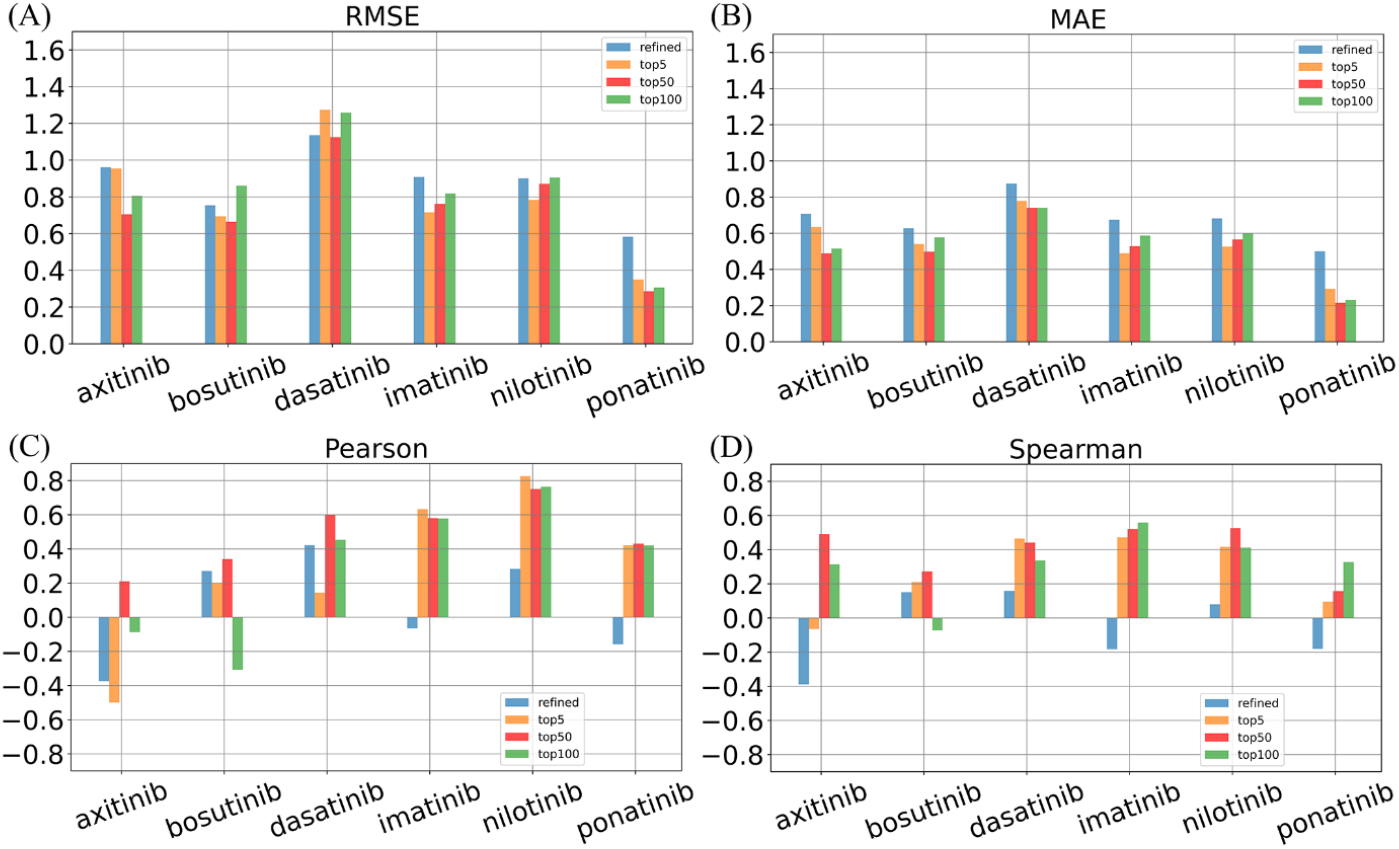
Evaluation metrics for using different number of generated conformational samples. Performance is evaluated using RMSE, MAE, Pearson coefficients, and Spearman coefficients for a refined set of PDBbind, and our method is evaluated by using 5, 50, and 100 samples.

### Data augmentation improves prediction accuracy

We assess the impact of sample size on prediction performance by expanding the dataset to 5, 50, and 100 times the original dataset. As can be seen from **Fig. 8**, the prediction based on the original dataset results in higher errors and lower correlations. The original dataset fails to reflect the impact of mutations and conformation on free energy changes. The best performance is achieved when the number of data is expanded by 50-fold (denoted as top50) on the original dataset. The top50 dataset reduced the RMSE and MAE by the data enhancement strategy, especially in the case of bosutinib, dasatinib, and ponatinib. In contrast, top5 and top100 performed slightly lower than top50. This indicates that the proper choice in the number of selected conformations can further improve the accuracy of the model prediction. From Pearson and Spearman coefficients, both top5 and top50 showed higher correlations on these datasets, especially top50 showed the best correlation on almost all drugs. The top50 also shows less variance over other datasets as shown in Fig. S10. The top50 may effectively achieve the balance between capturing the structural diversity and minimizing noise, while the top100 dataset may include excessive conformational changes, which also introduces variability in prediction. Due to the diverse conformational distribution among TKIs, both excessive and insufficient data can hinder improvements in model performance. The result also indicates that data augmentation strategies play an important role in improving model performance. Optimal performance will be achieved when conformational diversity is adequately represented without introducing redundancy or noise. Further optimizing the ratio of the number of samples can be conducted in the future.

### Conformations distribution indicates rational choice of conformations

The RMSD distribution characterizes the relative distribution of proteins between the active and inactive states after mutation. We characterize the filtered conformations of mutants in our model by clustering conformations based on their RMSD from the reference crystal structures. The clustering result in **Fig. 9A** reflects the preference of TKIs for kinase conformations. Cluster 1 is near the active state region, cluster 2 is near the inactive state region, and cluster 3 is in the middle transition region, which is validated by the extracted structures of each cluster center. Similar binding modes are observed for six TKI molecules as shown in **Fig. 9B** and Fig. S11. We also observe the changes in conformational distributions in the three clusters as shown in **Fig. 9C**. The selection process guided by the most probable distribution during data augmentation partially addresses the question of how to appropriately select a proper structure from ensembles. We demonstrate the application of subsampled AF2 in the downstream prediction of drug resistance. This approach offers new insights into studying the dynamic properties of protein-ligand interactions in drug discovery.

**Fig. 9:**
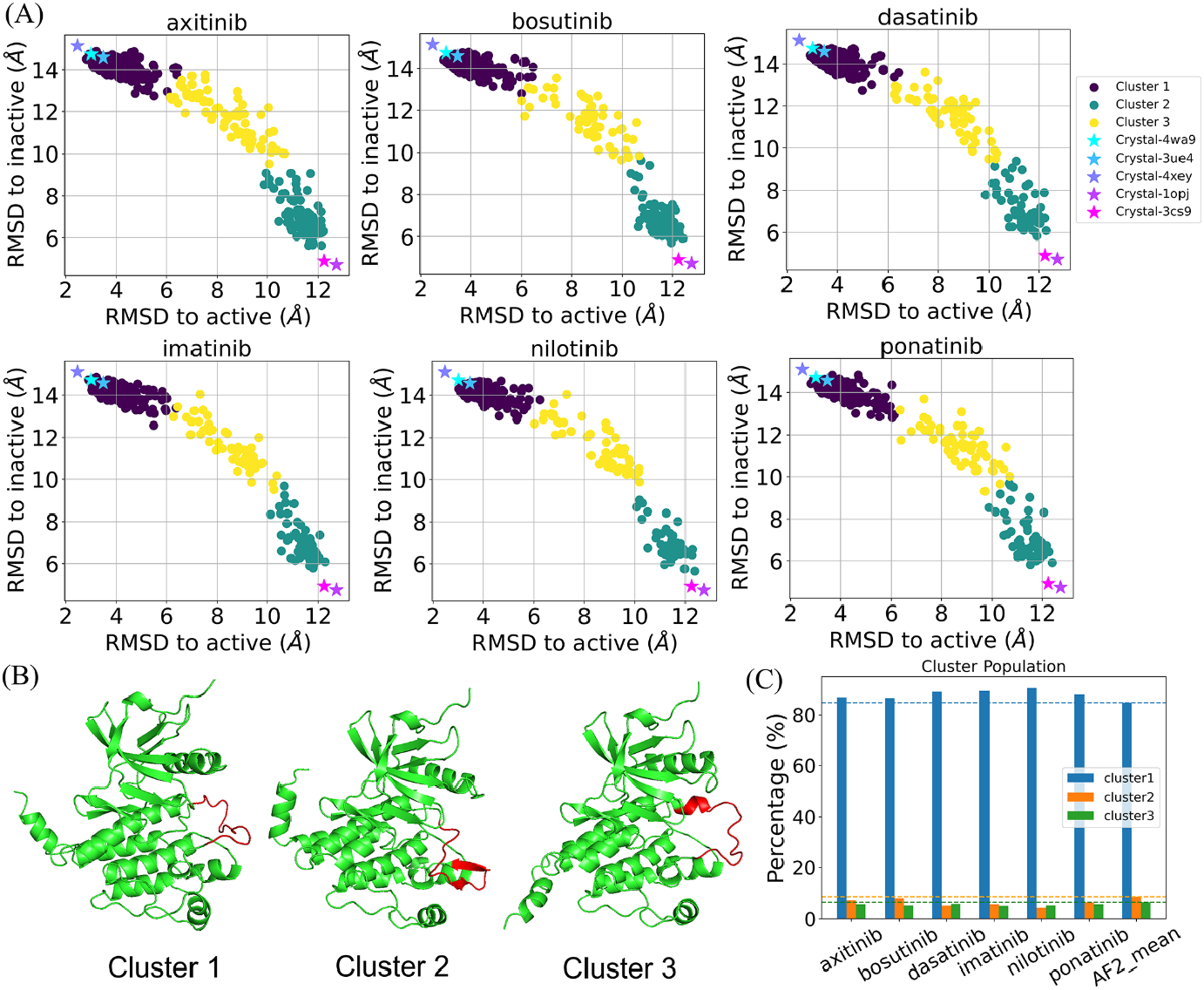
RMSD clustering of top50 dataset and the representative conformations. (A) RMSD distributions of conformations docked for six TKIs. (B) Representative conformations in each cluster center. (C) Percentage changes in each cluster after filtering. Crystal structures are labeled as the stars. The A-Loop region is highlighted in red in the structures.

## Conclusion

We present a method for predicting relative free energy in protein mutations by generating the relative distribution of conformations through AF2 subsampling. The data enhancement approach significantly improves the prediction performance of mutation effects by integrating the structural diversity of conformational ensembles, implementing a filtering strategy, and Siamese learning. Benchmarked using the TKI dataset and a refined set of PDBbind, the results demonstrate that the dataset augmented by 50-fold (top50), generated by using AF2 subsampling and Gaussian fitting, not only increases the structural diversity of mutation-ligand pairs but also effectively captures potential interaction patterns between different mutations and ligands. The STGNet model proposed in this study successfully captures the key features of protein-ligand interactions, thereby significantly improving the accuracy and stability of mutation effect predictions. We extend the application of integrating subsampled AF2 with Siamese learning to predict relative binding affinity changes, which holds promise for investigating drug resistance. This approach provides a promising tool for predicting the relative binding energy of mutation-ligand interactions without relying on the crystal structures of protein mutants.

## Methods

### Mathematical formulation

The energy difference between two states can be analyzed using perturbation theory. The perturbation depends on the selection of the reference state. In cases where the two states differ in the bound ligands, the perturbation corresponds to the energy difference in binding affinity induced by variations in the ligand. If two states differ in the conformational states of the protein, the perturbation reflects the energy difference in binding affinity resulting from conformational changes, which can be expressed as follows.

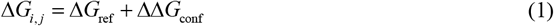

Transitioning from absolute binding energy Δ*G*_*i, j*_ to relative binding energy ΔΔ*G*_*i, j*_, we can define experimental value as the reference state, and the perturbed state is represented by the relative binding energy obtained from molecular docking. Thus, in our case, we can interpret the distribution of relative docking scores as the perturbations applied to the experimental distribution. The energy difference is computed from the reference states plus the energy variation as follows.

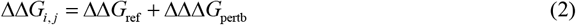

Therefore, the purpose of employing deep learning is to compute the differences in relative binding energy ΔΔΔ*G*_pertb_ . After generating the conformation ensemble, we conduct the molecular docking. We obtain the relative docking score difference between mutants and the wild-type protein.

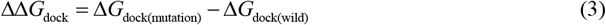

The relative docking score is used to select the most probable distribution based on the deviation from the Gaussian fitting mean value. The selected docking score and their corresponding conformation will then be paired with the experimental values. In the current study, we select the experimental value as the reference state. The difference in relative binding energy is computed from the following equation:

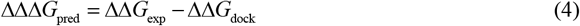

Here, ΔΔ*G*_exp_ is the energy of reference state, ΔΔΔ*G*_pred_ represents the energy difference caused by the conformation change and ligand variety. We aim to include conformational diversity in the prediction. Siamese networks are particularly well-suited for learning the differences between the two states, so we constructed a relative difference based on the experimental values and generated values, which serves as the regression target for our network learning.

The molecular representations and the difference binding energies are input into STGNet. After training the model, we predicted the binding energy difference ΔΔΔ*G*_pred_ for the test dataset. Siamese network was employed to learn the energy differences resulting from both conformation changes and mutations in protein sequences. The predicted energy differences ΔΔΔ*G*_pred_ are recovered to the predicted relative binding energy ΔΔ*G*_pred_ from the following equation:

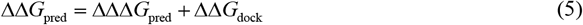

Each experimental value has been paired with N-generated samples, thereby we obtained N predicted relative binding energy. The final predicted relative binding energy is the average of the predicted relative binding energy:

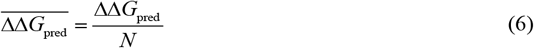

From the above workflow, we introduce the conformation ensemble in the prediction of relative binding energy. The relative binding energy can be recalibrated to the experimental level 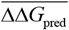 by computing the statistical average of STGNet predictions for the specified ligand molecule.

### Workflow for constructing the augmented dataset

As shown in **Fig. 2**, the workflow outlines a systematic approach to enhancing drug docking studies through the integration of reference data, conformational generation, and statistical analysis. By utilizing AlphaFold2 for conformational modeling and implementing ensemble docking procedures, we obtain relative docking scores that are subsequently analyzed using Gaussian fitting. The workflow emphasizes ranking these scores, filtering conformations based on probability proximity to the average, and ultimately constructing an augmented dataset.

#### Step 1: Construct reference data

Compile the original dataset, which includes 31 types of mutations and 6 associated compounds. Prepare mutation-ligand pairs and the corresponding relative binding energies.

#### Step 2: Generate conformations and conduct the docking

Utilize subsampled AlphaFold2 to generate protein conformations for wild and 31 types of ABL mutants. We generate 640 conformations for each type of mutation. Conduct molecular docking simulations on these generated conformations to assess binding affinities.

#### Step 3: Obtain docking scores and perform Gaussian fitting

Calculate the docking scores from the docking using AutoDock Vina. Check Gaussian fitting on the distribution of these docking scores to identify statistical averages.

#### Step 4: Prepare relative docking scores

Obtain relative docking scores between the wild and mutants to evaluate the binding efficacy of each conformation.

#### Step 5: Filter conformations

Filter the conformations and their corresponding relative docking scores to identify those that are close to the average score with the highest probability density.

#### Step 6: Construct an augmented dataset

Create an augmented dataset that includes the filtered conformations, their associated relative docking scores, and the difference in the relative docking scores. Ensure that the augmented dataset captures variability and distinct conformational states for further analysis.

### Datasets and preprocessing

#### TKI dataset

We validated the proposed approach on the TKI dataset collected by Hauser et al., which includes binding affinity data for mutation-ligand pairs (ΔΔ*G* in Kcal/mol). The ‘nd’ values were excluded based on the original dataset, resulting in the selection of 144 valid docking energy data for the mutants. In total, 31 mutant and 1 wild-type proteins were selected. The conformations from each protein were then adopted as the target to dock with different TKI molecules using AutoDock Vina, thus obtaining docking energy data for both mutant and wild-type conformations. The investigated TKI molecules include axitinib, bosutinib, dasatinib, imatinib, nilotinib, and ponatinib. The data is partitioned into training and test sets for cross-validation based on the name of TKI.

### Conformational ensemble prediction from AF2 subsampling

Monteiro et al. developed subsampled AlphaFold2 to predict the relative populations of protein conformations by subsampling MSA. Their results show that the subsampling method can accurately predict the relative conformational populations of proteins when compared to the data from NMR experiments. They also obtained the accurate relative state population changes for the eight mutants for ABL kinase. We employed the same implementation to sample conformation ensembles of 31 mutants of ABL kinase. The residue length indices from 248 to 534 according to the crystal structure (PDB ID: 6XR6), following the same prediction procedure using ColabFold. ColabFold utilizes the fast homology search of MMseqs2 with AF2 to facilitate rapid prediction of protein structures and complexes. The value of max_seq:extra_seq was set to 256:512, which would result in the most diverse results concerning the activation loop conformation. We utilized 5 models of AF2 per prediction, performed 4 rounds of optimization, and set the number of seeds as 32. In total, we generated 640 conformations for each mutation.

### Molecular docking and preprocessing of the relative binding energy

Different conformations of the proteins were obtained through subsampling, resulting in a relative conformational distribution. Molecular docking for each sampled conformation was performed using AutoDock Vina to determine the binding energy of the ligand across different mutations as shown in Step 2 in **Fig. 2**. The docking center is set as the coordinate center of four residues with residue index 267, 288, 378, and 399. The exhaustiveness of AutoDock Vina is set as 32. The docking energies for both the mutant and wild type were derived from ensemble docking. We computed the difference in docking score between the mutant and the wild type in Step 3 and Step 4. Then we filtered the conformational data with the smallest deviation from the Gaussian mean of the binding energy distribution of each mutant-ligand combination in Step 5. We selected the top *k* (eg. 1, 5, 50, and 100) values with the smallest absolute differences value from the Gaussian mean value for pairing. The *k* equals 1, which means no flexibiligy is incuded. We select the *k* values near the average because the Gaussian fitting indicates that values close to the average have the highest probability distribution. We calculate ΔΔΔ*G*_cal_ different TKIs by calculating the differences between the experimental value and the generated value in Step 6. Finally, all composite confirmation data are grouped to construct the required data-augmented dataset. The total number of raw predictions is equal to the number of conformations multiplied by the number of ligands, represented as *N*=*k*×6. As shown in **Fig. 2**, conf_exp consists of a single conformation, while conf_AF comprises *k* conformations generated from subsampled AF2 corresponding to the same mutation type.

### Refined set of PDBbind for direct absolute value prediction

The refined set of PDBbind version 2020 was selected to compare with our approach. After the removal of ABL samples, the refined set was divided into the training and validation sets in a 9:1 ratio. The CASF-2016 is used to benchmark the performance of the regular network model. The test set data comprises the 144 mutation data in the original TKI dataset alongside the six TKIs for wild type. The training was conducted using a similar network architecture, except for removing the Siamese network. After training the model, we predict the absolute binding free energy separately for both the wild type and mutations. Subsequently, we computed the relative binding affinity by subtracting the absolute binding energy of mutation from that of the wild type, respectively.

### Input features of the Siamese learning

We adopt both the structural information and molecular graph as the molecular representation to capture the contribution from the conformation and the interaction between proteins and ligands. The molecular representations are summarized in Table S6.

### Protein contact maps

Predicting geometrical relationships between residues (including distances and orientations) has demonstrated its importance in structure predictions. Protein contact maps use the adjacency matrix *A*_*i*, *j*_ to represent the graph structure of a protein, which contains information about whether or not amino acids are found in the protein chain, specifically, a protein map *G* = (*N*_*p*_, *M* _*p*_) is a contact map generated by taking an input protein containing *L*_*p*_ residues. A pair of residues is considered to be in contact when the Euclidean distance *d*_*i, j*_ between the CA atoms of the two pairs of amino acids is less than a threshold *d*_*c*_ . The residues in a pair are considered to be in contact with each other. These connections are determined from the three-dimensional structure of the proteins. We set a threshold of 10 Å.

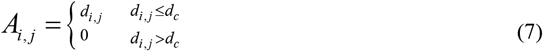

The protein contact map of the MPNN structure can be represented as *G*_*contactMap*_ = (*V*, *E*), where *V* denotes a set of *n* atomic nodes, and each node feature vector denotes (*x*_*v*_), including the one-hot coding of the protein sequence, the 12 amino acid nature features, and the BLOSUM62 matrix features^62^. *E* denotes the relationship of the edges of the neighbor matrix in the molecular graph, and the eigenvector of each edge is denoted (*e*_*vw*_) as the Euclidean distance between each pair of amino acids.

### Ligand graph

We choose the molecular graph as the representation of the ligands. The molecular graph of the ligand for the MPNN structure can be represented as *G*_*ligand*_ = (*V*, *E*) . where *V* denotes a set of n atomic nodes, each represented by an eigenvector of 75 values (*x*_*v*_), including 44 atomic types, 11 atomic degrees, 7 implicit valence states, charge, number of free electrons, 5 hybridization types, aromaticity, and 5 hydrogen atoms. *E* represents the relationship of the edges of the neighbor matrix in the molecular graph. Each edge is represented by 6 values called eigenvectors (*e*_*vw*_), including 4 bond types, conjugacy, and intra-ring bonds. The molecular graph has been validated in our previous work.

### Protein-ligand interaction graph

We adopt the 3D interaction graph structure data used by GIGN. The complex is represented as a 3D interaction graph *G*_*inter*_ = (*V*,ε, *R*) = (*V*_*l*_ ⏝*V*_*p*_,ε_*l*_ ⏝ε_*lp*_, *R*_*l*_ ⏝ *R*_*p*_), where *V*_*l*_ and *V*_*p*_ are sets of ligand and protein nodes (atoms), ε_*l*_ and ε _*p*_ are sets of ligand and protein covalent edges (covalent bonds/covalent interactions). *R*_*l*_ and *R*_*p*_ are sets of ligand and protein 3D coordinates. Each node *v*_*i*_ ∈*V* has initial features *x* _*i*_ ∈ R ^*n*^ and 3D coordinates *r* ∈R^3^ . When atoms *v*_*i*_ and *v* _*j*_ are connected by a covalent edge, the covalent edge *e*_*ij*_ ∈ε_*l*_ ⏝ε _*p*_ . A noncovalent edge *e* ∈ε exists when *d* < 15 Å, where *d*_*ik*_ = ‖ *r* _*i*_ − *r* _*k*_ ^2^ ‖ is the distance between the ligand atom _*ik lp*_ *v*_*i*_ ∈*V*_*l ik*_ and the protein atom *v*_*k*_ ∈*V*_*p*_ .

### Siamese learning

#### Message Passing Neural Networks

We used message-passing networks to process contact maps for proteins and graph data for ligands. Gilmer et al. designed an MPNN framework for processing chemical graphs. Forward propagation consists of two phases, the message-passing phase and the readout phase. The message passing phase runs for T time steps and is defined according to the message function *M*_*t*_ and vertex update function *U*_t_ . In the message passing phase, the hidden state 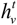 of each node is updated according to the message 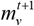 as follows.

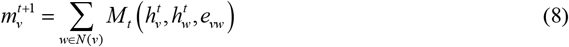

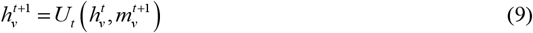

Where, *N* (*v*) denotes the neighbor of node *v* in graph *G*, *M*_*t*_ denotes the message function, *U*_*t*_ denotes the vertex update function. The readout phase uses the readout function *R* to compute the feature vectors of the entire graph according to the following equation.

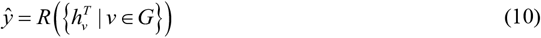

where *R* denotes the readout function.

### Geometric interaction graph neural network

Yang et al. designed an ensemble interaction graph neural network, a Geometric interaction graph neural network (GIGN) that combines 3D structural and physical interactions to learn node representations more efficiently through a heterogeneous interaction layer that unifies covalent and non-covalent interactions into a message-passing phase.

In the Geometric Interaction Graph Neural Network (GIGN), the process of integrating covalent and noncovalent interactions into node representations is handled through a message-passing mechanism that accounts for the heterogeneity of these interactions. Each node in the 3D interaction graph, representing an atom, updates its representation by aggregating messages from its neighboring nodes, which are categorized as either covalent or noncovalent interactions. The messages from covalent neighbors are calculated using a function *M* (_cov_) that incorporates the node features *x* and their 3D coordinates *r*, while messages from noncovalent neighbors are computed using a similar function *M* ^(ncov)^ . These messages are then aggregated separately, respecting their distinct nature.

The aggregated messages, which differ in scale due to the significant difference in the number of noncovalent versus covalent interactions, are normalized to the same scale by multilayer perceptrons *U* (_cov_) and *U* (_ncov_) before integration. This integration step ensures that the information from both interaction types is effectively combined to refine the node’s representation. The final representation update for each node is given as follows.

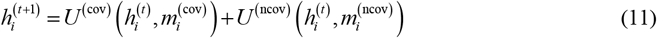

Where, 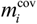 and 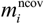 are the aggregated messages from covalent and noncovalent neighbors, respectively, and 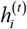 is the current representation of node *v*_*i*_ at layer *t* . This approach maintains the invariance property concerning input translations and rotations, avoiding the need for computationally expensive data augmentation and leading to a more efficient and biologically meaningful prediction of protein-ligand binding affinities.

### STGNet model

We propose the SGTNet model for predicting the relative binding energy for protein-ligand complexes. The SGTNet model uses dual inputs of experimental data and generated data as shown in **Fig. 3**. The pairings are made between the reference conformation and the generated conformations. This enables the fusion of ligand-binding features with information on the structural changes triggered by the mutation. For the learning with protein contact maps and small molecules, we take the graph information as inputs and perform feature extraction through the MPNN model with shared parameters to get the feature differences between the two inputs, followed by further extraction of the deep features through the fully connected layer. For the learning of protein-ligand interactions, we take the graph information of the docked protein-ligand structure as inputs perform feature extraction via the GIGN model with shared parameters, and further extract deep features via the fully-connected layer after calculating the feature differences. Then, all the learned graph features are concatenated together to form a joint feature, which generates the final prediction ΔΔΔ*G*_pred_ through a fully connected layer.

To obtain the target prediction value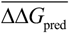, the calculation method is shown in Fig. S1. The results of the raw predictions for a given test set include specific ligands, and each mutation corresponds to the predicted binding energies for these ligands. Consequently, the target prediction value needs to be averaged from the raw predicted results. We evaluated the STGNet model using the original dataset, and three pairwise augmented datasets with *k* chosen as 5, 50, and 100, respectively.

## Author contributions

L.Xie.: Initiated the project and proposed the method. X.L.: Performed modeling. L.Xie. and X.L.: Conducted data analysis. D.Z. Y.J. and L.Xu.: Contributed the data analysis. S.C. and X.X.: Supervised the work.

## Data availability

AlphaFold2 subsampling prediction data generated in this study have been deposited in the Zenodo: https://zenodo.org/records/14607966.

## Code availability

The code used to generate the results shown in this study is available under an MIT License via GitHub at https://github.com/AIMedDrug/STGNet.

## Additional information

The detailed analysis can be found in the supplementary material.

## Key Points

- Predicting the impact of mutations on protein-ligand binding affinity is vital for addressing drug resistance and repurposing drugs. Current methods face challenges due to reliance on known protein-ligand structures.
- We proposed a novel few-shot learning framework that integrates AlphaFold2 subsampling, ensemble docking, and Siamese learning to predict relative binding affinity changes caused by mutations.
- We augmented the dataset by pairing the reference state with the structures generated by AlphaFold2. By incorporating the generated data, we enhance the model’s prediction accuracy and robustness, particularly for targets with high structural variability.
- The proposed method achieves higher prediction accuracy for relative binding affinities by incorporating structural diversity from AlphaFold.

