## Supplementary material for "Predicting Mutation-Induced Relative Protein-Ligand Binding Affinity Changes via Conformational Sampling and Diversity Integration with Subsampled Alphafold2 in Few-Shot Learning": Supplemetal Figure

### Supplementary Information

#### Supplemetary Figures

##### Calculation of free energy prediction values for original, top5, top50, and top100 for data augmentation methods

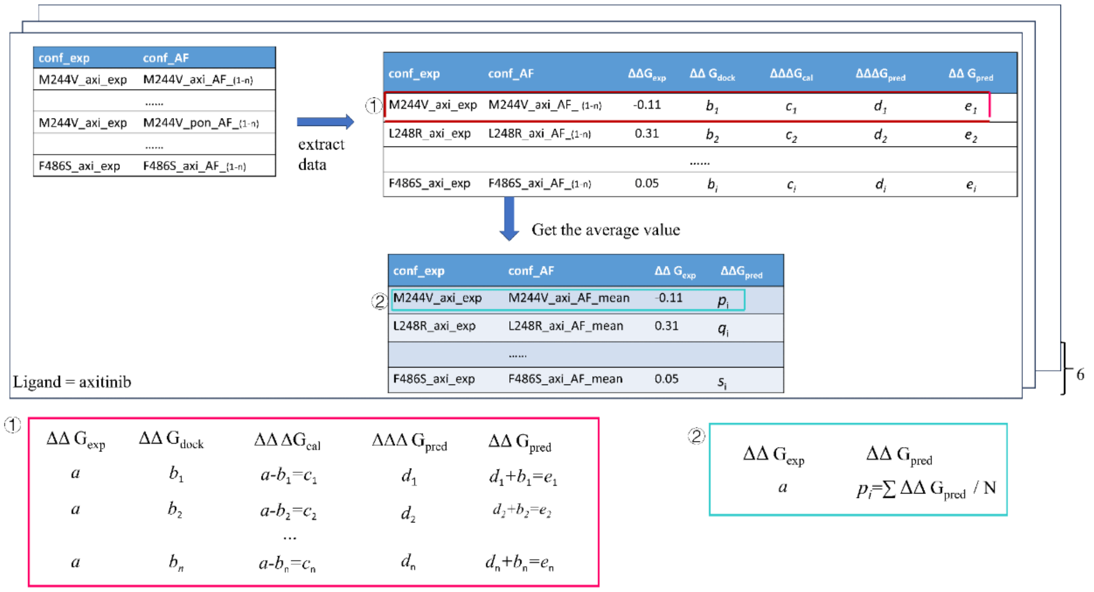


**Fig. S1** Calculation of free energy prediction values for the original dataset, top5, top50, and top100 for data augmentation methods.

Fig. S1 depicts the computational process $\Delta\Delta G_{\text{pred}}$. The conf_exp labeling of the same mutant-ligand corresponds to the predicted values of multiple conf_AF different conformations, and $\Delta\Delta G_{\text{pred}} = \Delta\Delta\Delta G_{\text{pred}} + \Delta\Delta G_{\text{dock}}$ eliminates errors in the docking data. The prediction $\Delta\Delta G_{\text{pred}}$ averaged over multiple conf_AF corresponds to the relative binding energy for the same mutation and ligand. The average value $\overline{\Delta\Delta G_{\text{pred}}}$ is the final relative binding energy, and is used for the performance evaluation of the model. The Pearson and Spearman coefficient is calculated between $\overline{\Delta\Delta G_{\text{pred}}}$ and the experimental value $\Delta\Delta G_{\text{exp}}$.

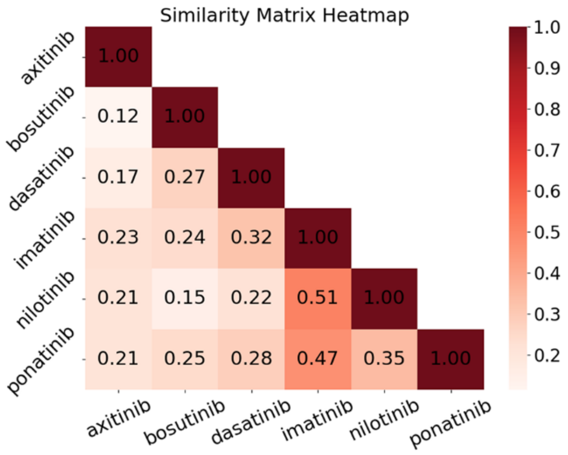


| TKIs | Types |
|---|---|
| **axitinib** | IIA |
| **bosutinib** | IIB |
| **dasatinib** | I/I½A |
| **imatinib** | IIA |
| **nilotinib** | IIA |
| **ponatinib** | I |

**Fig. S2** Tanimoto similarity map for the investigated TKI molecules. The left side shows the similarity plot of the TKI molecules. The right side indicates the types of different TKIs.

Fig. S2 shows the similarity graph for our investigated molecules. Each point on the map represents a pair of molecules, with darker colors indicating higher similarity and lighter colors indicating greater dissimilarity. These inhibitors work by competing with ATP for the ATP-binding site of the kinase, thereby inhibiting the kinase's activity by binding at different sites. All molecules exhibit low similarity; therefore, constructing pairs using ligand variation alongside the experimental values does not lead to information leakage.

#### Relative binding free energy distribution

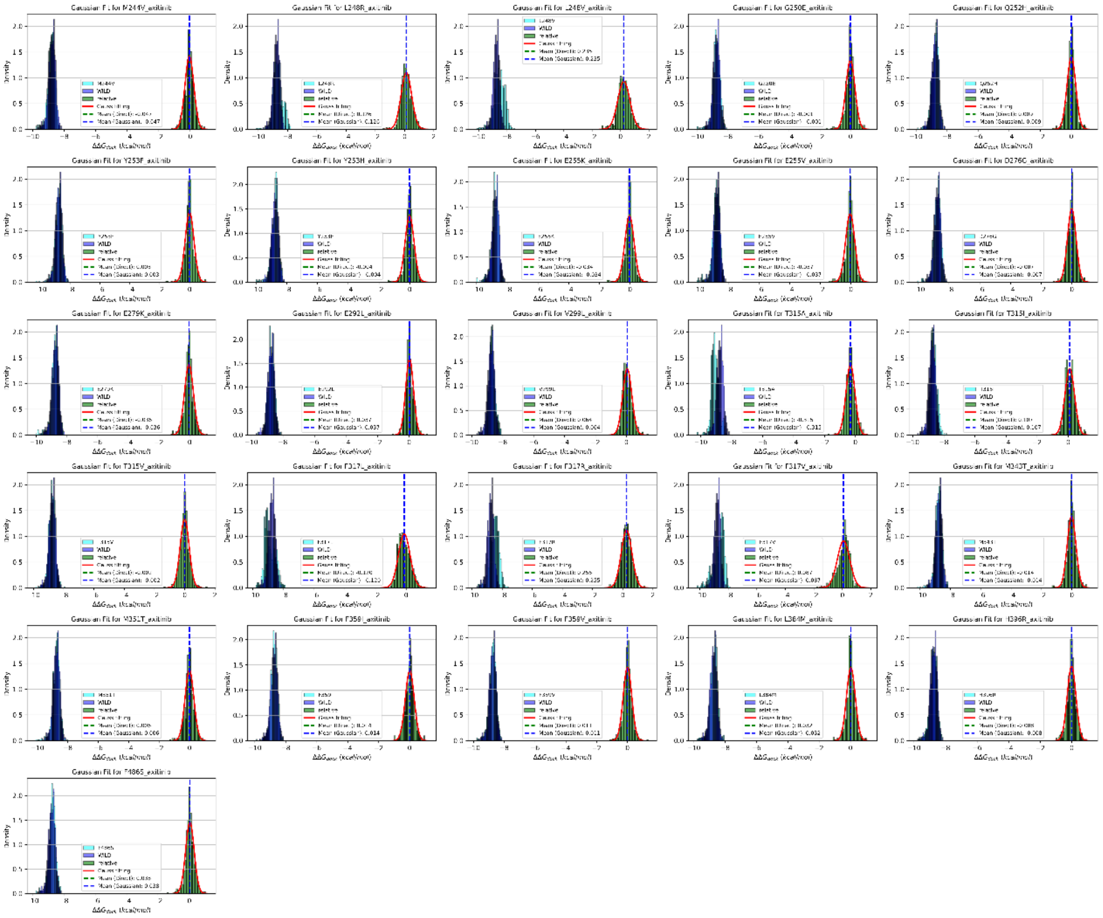

**Fig. S3** Distribution of docking score and the relative docking score for axitinib.

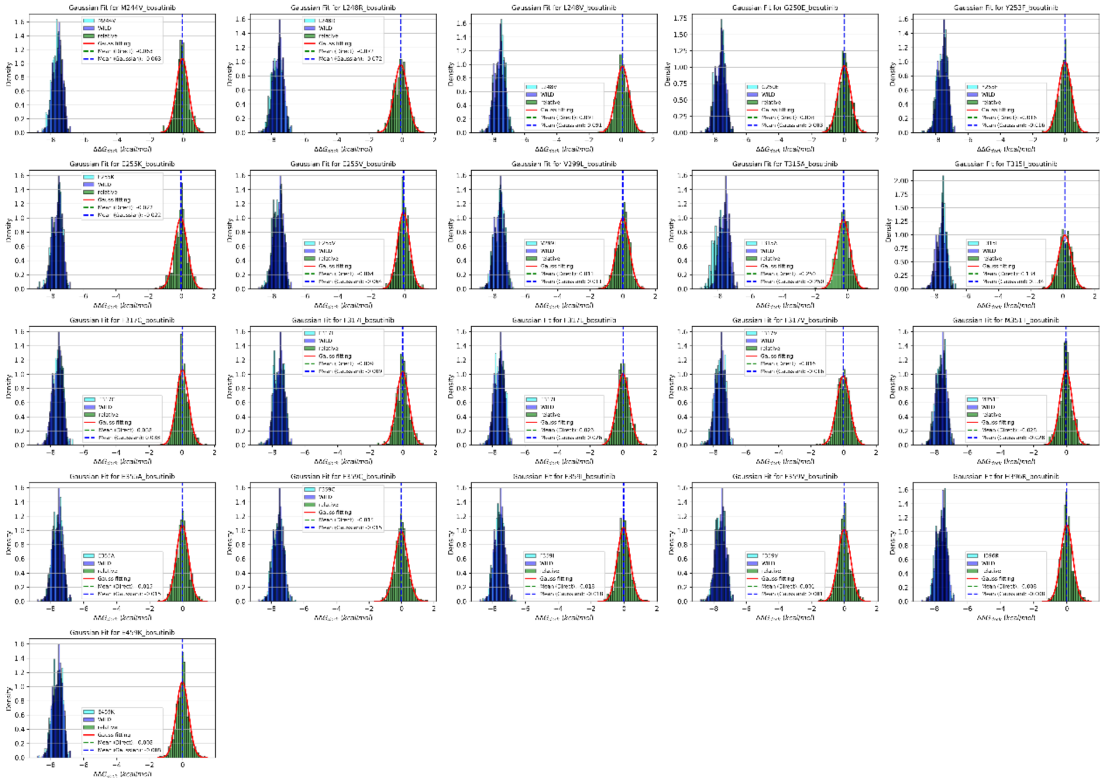

**Fig. S4** Distribution of docking score and the relative docking score for bosutinib.

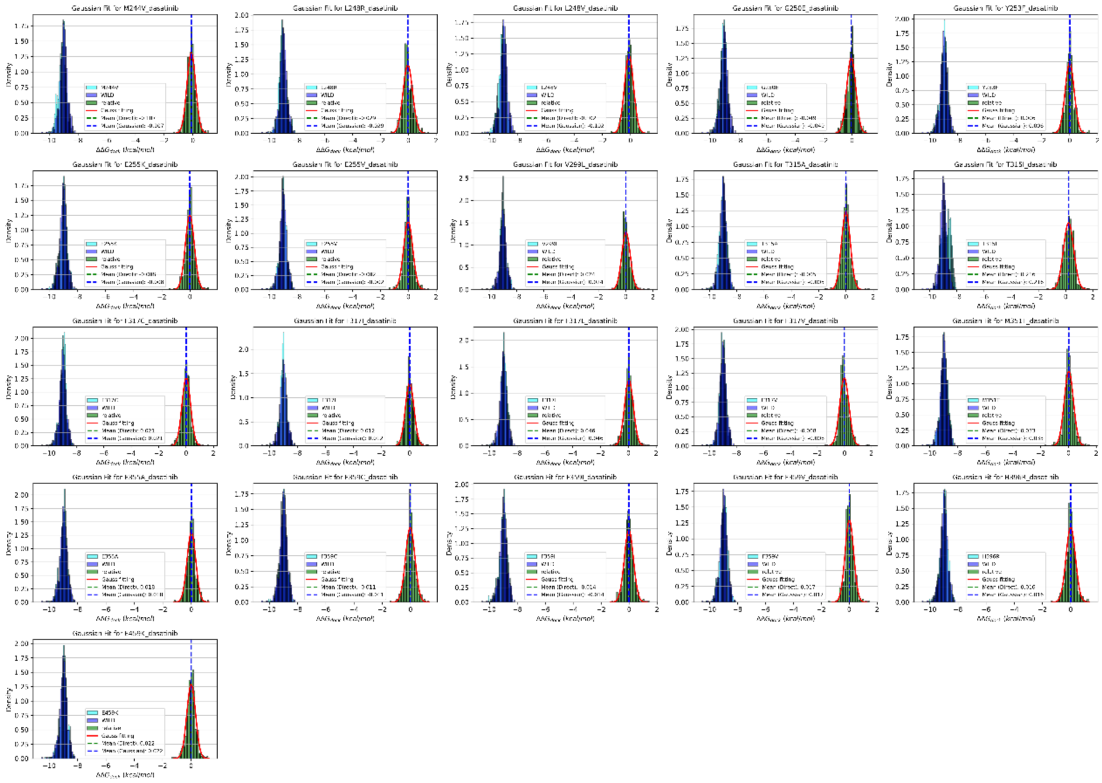

**Fig. S5** Distribution of docking score and the relative docking score for dasatinib.

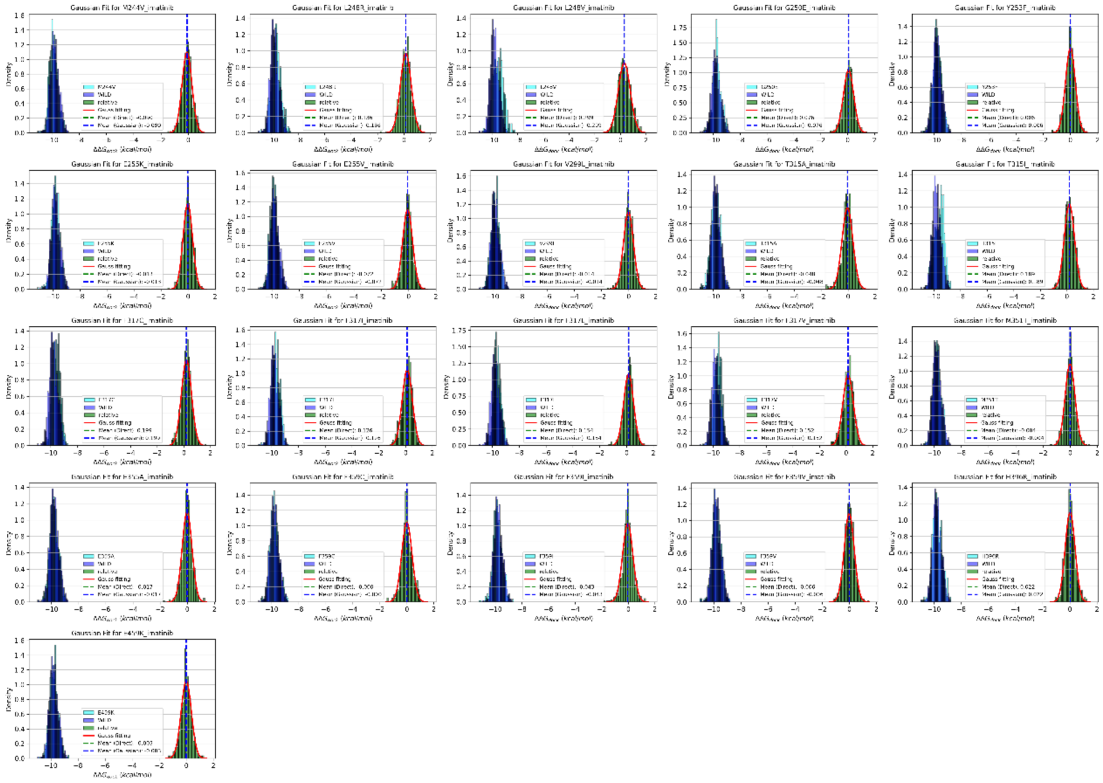

**Fig. S6** Distribution of docking score and the relative docking score for imatinib.

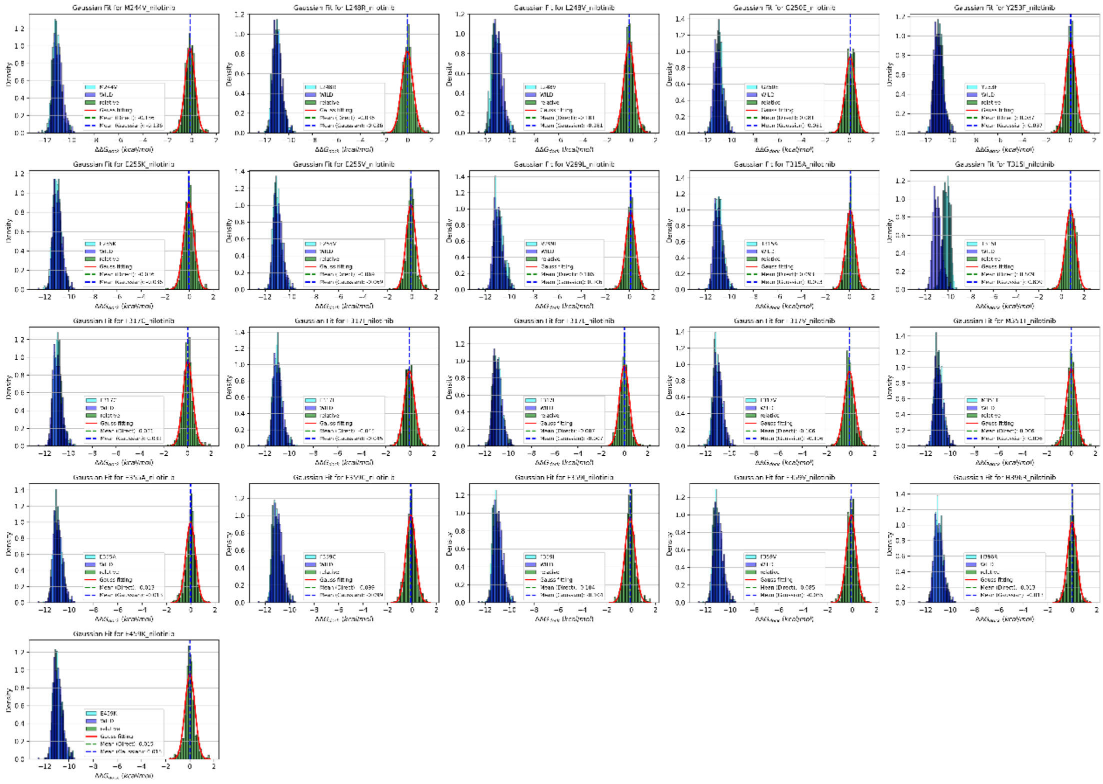

**Fig. S7** Distribution of docking score and the relative docking score for nilotinib.

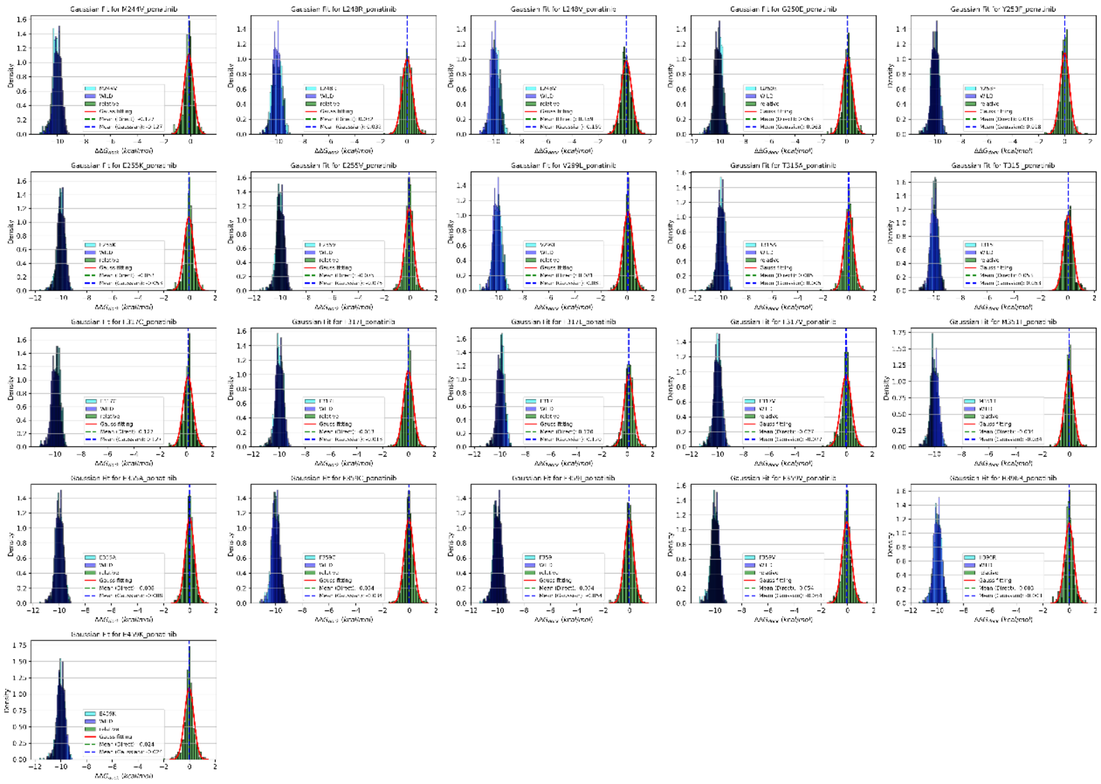

**Fig. S8** Distribution of docking score and the relative docking score for ponatinib.

##### Direct predicting absolute binding free energies using the refined set of PDBbind

Direct predicting absolute binding free energy adopts similar neural networks and the arichitecture is shown in Fig S9. Neural networks with multilevel graph representations learn protein contact maps and ligands using MPNN, protein-ligand interactions using GIGN, and concatenate the learned features to make predictions using a fully connected layer.

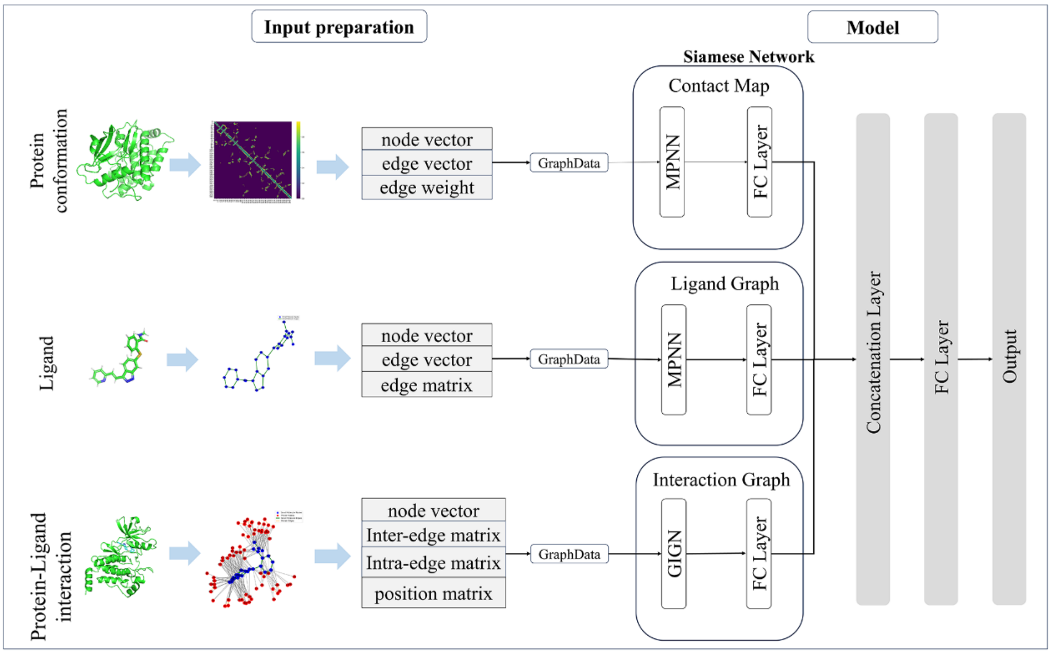


**Fig. S9** Neural network for processing three graph representations of refined datasets.

To evaluate the stability and reliability of the results, five repeated calculations were performed using the direct absolute prediction based on the refined set of PDBbind and the data augmentation based on the top5 and top50 datasets. The variances of RMSE, MAE, Pearson coefficients, and Spearman coefficients for different models were calculated and shown in Fig. S10.

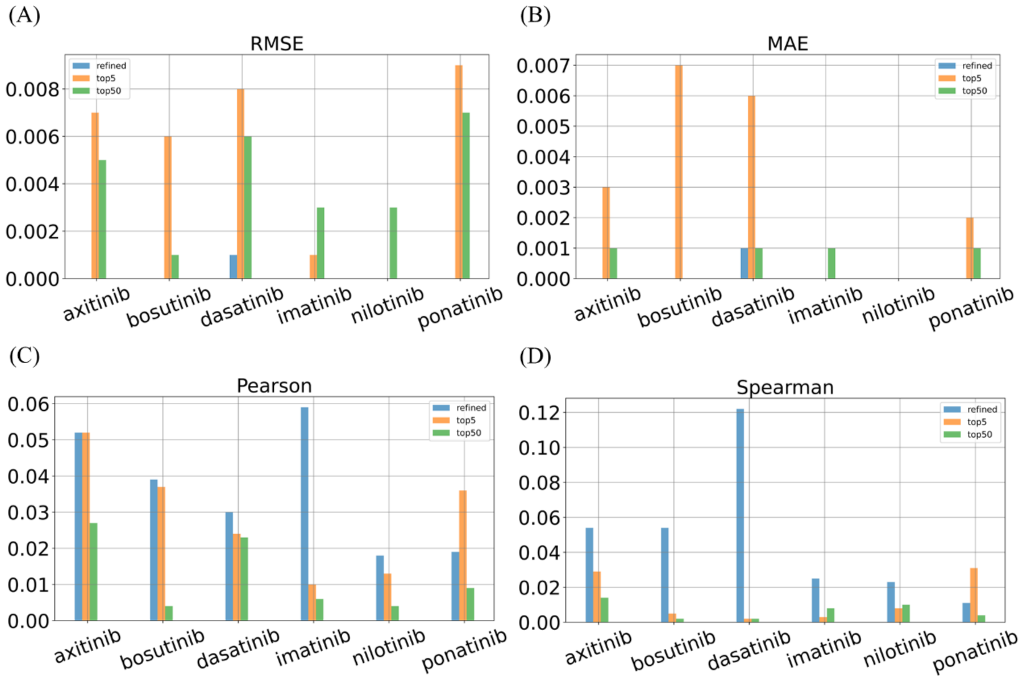


**Fig. S10** Variance of RMSE, MAE, Pearson coefficients, and Spearman coefficients in five rounds of computation.

**Representative conformations in three clusters for different TKIs**

Figure S11 shows representative conformations of various tyrosine kinase inhibitors (TKIs), divided into three different clusters. Among them, cluster 1 is located around the active state, cluster 2 is located near the inactive state, and cluster 3 is the intermediate state. In cluster 1, the individual TKI representative structures exhibit active state features, including the openness of the kinase active site and the active conformation of the activation loop, which is consistent with the structural properties required for ATP binding or kinase activation. In cluster 2, the individual TKI representative structures exhibit significant features of the inactive state, exhibiting a steady state of inhibition upon self-inhibition or binding of an inhibitor. In cluster 3, the individual TKI representative structures are characterized by a dynamic intermediate state in which the openness of the active site and the compactness of the structural domains are balanced with each other.

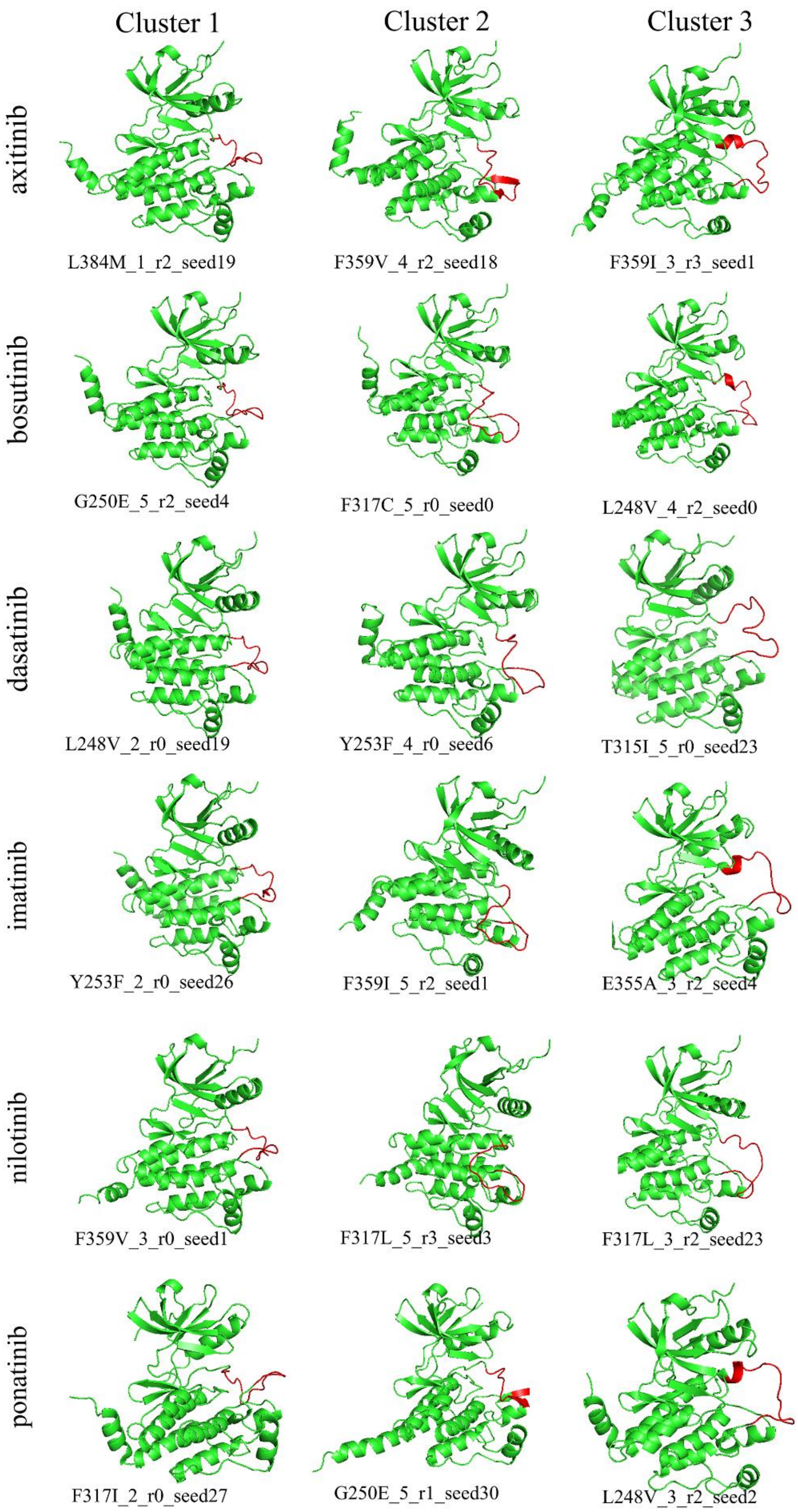


**Fig. S11** Representative conformations in three clusters.

#### Supplemetary Tables

#### Relative population of conformations

**Table S1**. The population distribution of conformation ensemble of ABL kinase

| Protein | Ground state population | Ratio of cluster 1 | Ratio of cluster 2 | Ratio of cluster 3 |
|---|---|---|---|---|
| WILD | 83.0% | 84.2% | 10.2% | 5.6% |
| M244V | 84.5% | 85.5% | 9.8% | 4.7% |
| L248R | 79.8% | 81.6% | 7.3% | 11.1% |
| L248V | 83.8% | 84.7% | 5.3% | 10.0% |
| G250E | 82.2% | 84.2% | 9.5% | 6.3% |
| Q252H | 84.5% | 85.6% | 10.0% | 4.4% |
| Y253F | 80.6% | 81.7% | 9.5% | 8.8% |
| Y253H | 87.2% | 89.1% | 6.1% | 4.8% |
| E255K | 82.3% | 84.2% | 8.3% | 7.5% |
| E255V | 82.2% | 83.9% | 6.9% | 9.2% |
| D276G | 83.9% | 84.7% | 10.2% | 5.1% |
| E279K | 80.2% | 81.6% | 8.8% | 9.6% |
| E292L | 89.1% | 89.2% | 8.6% | 2.2% |
| V299L | 84.1% | 85.0% | 10.2% | 4.8% |
| T315A | 83.0% | 85.0% | 9.1% | 5.9% |
| T315I | 87.7% | 88.0% | 8.1% | 3.9% |
| T315V | 82.8% | 84.2% | 9.5% | 6.3% |
| F317C | 82.3% | 83.0% | 10.5% | 6.5% |
| F317I | 82.3% | 83.1% | 8.9% | 8.0% |
| F317L | 83.6% | 84.4% | 8.6% | 7.0% |
| F317R | 85.3% | 85.5% | 8.4% | 6.1% |
| F317V | 80.0% | 81.6% | 8.6% | 9.8% |
| M343T | 81.4% | 82.7% | 9.1% | 8.2% |
| M351T | 83.9% | 85.3% | 8.4% | 6.3% |
| E355A | 84.8% | 85.9% | 6.7% | 7.4% |
| F359I | 72.7% | 75.5% | 14.5% | 10.0% |
| F359V | 80.0% | 81.1% | 10.2% | 8.7% |
| L384M | 91.4% | 91.1% | 6.9% | 2.0% |
| H396R | 85.3% | 87.7% | 8.4% | 3.9% |
| F486S | 89.4% | 89.8% | 5.8% | 4.4% |
| E459K | 87.0% | 87.8% | 8.6% | 3.6% |

Table S1 presents the population distribution of conformation ensembles for ABL kinase. The table provides a detailed breakdown of the ground state population and the ratios of three distinct clusters for a series of mutations and ligand interactions. The ground state population represents the most stable conformation of the protein, while the ratios of cluster 1, cluster 2, and cluster 3 indicate the distribution of the protein's conformations within each cluster. This data is crucial for understanding the structural diversity of the protein and how it may affect its function and interaction with ligands.

#### Performance of RMSE, MAE, Pearson coefficients, and Spearman coefficients for different datasets with different TKIs

Molecular docking and MOESM5 are datasets constructed based on physics-based methods (molecular docking and FEP+). The refined set is a dataset constructed based on direct absolute value prediction. The original dataset, top5, and top50 are datasets constructed based on data enhancement methods. The evaluation metrics of these datasets are shown in Tables S2-S5. The best score is highlighted in red color.

**Table S2**. RMSE of the evaluated methods for different TKI

| RMSE | axitinib | bosutinib | dasatinib | imatinib | nilotinib | ponatinib |
|---|---|---|---|---|---|---|
| docking | 0.622 | 0.820 | 1.391 | 0.966 | 0.829 | 0.406 |
| refined | 0.961 | 0.754 | 1.136 | 0.908 | 0.901 | 0.583 |
| MOESM5 | 0.907 | 1.406 | 1.040 | 1.061 | 1.061 | 1.087 |
| original | 0.876 | 0.692 | 1.257 | 0.902 | 0.868 | 0.277 |
| top5-mean | 0.955 | 0.694 | 1.274 | 0.716 | 0.783 | 0.350 |
| top50-mean | 0.705 | 0.664 | 1.125 | 0.762 | 0.871 | 0.286 |

**Table S3**. MAE of the evaluated methods for different TKI

| MAE | axitinib | bosutinib | dasatinib | imatinib | nilotinib | ponatinib |
|---|---|---|---|---|---|---|
| docking | 0.387 | 0.597 | 0.966 | 0.719 | 0.594 | 0.330 |
| refined | 0.708 | 0.628 | 0.875 | 0.673 | 0.682 | 0.501 |
| MOESM5 | 0.705 | 0.960 | 0.734 | 0.782 | 0.815 | 0.867 |
| original | 0.688 | 0.488 | 0.781 | 0.590 | 0.568 | 0.224 |
| top5-mean | 0.635 | 0.540 | 0.779 | 0.490 | 0.526 | 0.293 |
| top50-mean | 0.489 | 0.498 | 0.741 | 0.528 | 0.566 | 0.215 |

**Table S4**. Pearson coefficient of the evaluated methods for different TKI

| Pearson | axitinib | bosutinib | dasatinib | imatinib | nilotinib | ponatinib |
|---|---|---|---|---|---|---|
| docking | -0.043 | 0.099 | 0.545 | -0.251 | 0.554 | 0.014 |
| refined | -0.375 | 0.271 | 0.423 | -0.065 | 0.283 | -0.158 |
| MOESM5 | 0.566 | -0.148 | 0.768 | 0.565 | 0.590 | 0.228 |
| original | -0.021 | 0.302 | 0.684 | 0.014 | 0.659 | -0.236 |
| top5-mean | -0.499 | 0.199 | 0.143 | 0.633 | 0.827 | 0.422 |
| top50-mean | 0.210 | 0.340 | 0.600 | 0.580 | 0.751 | 0.431 |

**Table S5**. Spearman coefficient of the evaluated methods for different TKI

| Spearman | axitinib | bosutinib | dasatinib | imatinib | nilotinib | ponatinib |
|---|---|---|---|---|---|---|
| docking | 0.106 | 0.188 | 0.262 | -0.149 | 0.373 | 0.089 |
| refined | -0.389 | 0.151 | 0.158 | -0.183 | 0.080 | -0.181 |
| MOESM5 | 0.454 | -0.157 | 0.429 | 0.298 | 0.220 | 0.213 |
| original | 0.1 | 0.103 | 0.212 | 0.111 | 0.081 | -0.265 |
| top5-mean | -0.065 | 0.210 | 0.465 | 0.472 | 0.417 | 0.096 |
| top50-mean | 0.492 | 0.273 | 0.441 | 0.521 | 0.526 | 0.157 |

#### Molecular representations

Molecular representations includes three parts: contact map for proteins, graph for ligands, and graph for protein-ligand interactions. The features are summarized in Table S6.

**Table S6**. Molecular representations for the STGNet.

| Type | Representation | Features |
|---|---|---|
| Structure | Contact map | 1. Node features: (1) One hot for residue names. (2) residue features: aliphatic (AILMV), aromatic(FWY), polar neutral (CNQST), acidic charged(DE), basic charged(HKR), weight, pka, pkb, logarithm of the dissociation constant, isoelectric point of residues, hydrophobicity of residue at pH2 and pH7. (3) BLOSUM62 vectors.<br>2. Edge weights: distance of contacted CA atom devided by 10 |
| Graph | Ligand | 1. Atom features: atom symbol, atom degree, atom implicit valence, atomic formal charge, number of radical electrons, atom hybridization, aromatic.<br>2. Bond features: bond type (single, double, triple, aromatic), bond conjugation, bond in ring |
| | Protein Ligand Interaction | 1. Protein atoms: atom_symbols, atom degree, atom hybridization, aromatic, number of hydrogens.<br>2. Ligand atoms: atom symbol, atom degree, atom implicit valence, atomic formal charge, number of radical electrons, atom hybridization, aromatic, number of hydrogens.<br>3. Edge: add edges when distance_threshold=15 |
